# Local assembly of long reads enables phylogenomics of transposable elements in a polyploid cell line

**DOI:** 10.1101/2022.01.04.471818

**Authors:** Shunhua Han, Guilherme B. Dias, Preston J. Basting, Raghuvir Viswanatha, Norbert Perrimon, Casey M. Bergman

## Abstract

Animal cell lines cultured for extended periods often undergo extreme genome restructuring events, including polyploidy and segmental aneuploidy that can impede *de novo* whole-genome assembly (WGA). In *Drosophila*, many established cell lines also exhibit massive proliferation of transposable elements (TEs) relative to wild-type flies. To better understand the role of transposition during long-term animal somatic cell culture, we sequenced the genome of the tetraploid *Drosophila* S2R+ cell line using long-read and linked-read technologies. Relative to comparable data from inbred whole flies, WGAs for S2R+ were highly fragmented and generated variable estimates of TE content across sequencing and assembly technologies. We therefore developed a novel WGA-independent bioinformatics method called “TELR” that identifies, locally assembles, and estimates allele frequency of TEs from long-read sequence data (https://github.com/bergmanlab/telr). Application of TELR to a ∼130x PacBio dataset for S2R+ revealed many haplotype-specific TE insertions that arose by somatic transposition in cell culture after initial cell line establishment and subsequent tetraploidization. Local assemblies from TELR also allowed phylogenetic analysis of paralogous TE copies within the S2R+ genome, which revealed that proliferation of different TE families during cell line evolution *in vitro* can be driven by single or multiple source lineages. Our work provides a model for the analysis of TEs in complex heterozygous or polyploid genomes that are not amenable to WGA and yields new insights into the mechanisms of genome evolution in animal cell culture.

## Introduction

Cell lines are commonly used in biological and biomedical research, however little is known about how cell line genomes evolve *in vitro*. For decades, it has been well-established that immortalized cell lines derived from plant or animal tissues often develop polyploidy or aneuploidy during routine cell culture (Ford and Yerganian 1958; Hink 1976; Ogura 1990; Bairu *et al*. 2011). More recently, the use of DNA sequencing has further revealed that segmental aneuploidy and other types of submicroscopic structural variation are widespread in cell lines (Zhang *et al*. 2010; Miyao *et al*. 2012; Adey *et al*. 2013; Lee *et al*. 2014; Nattestad *et al*. 2018; Ben-David *et al*. 2018; Zhou *et al*. 2019b,a; Liu *et al*. 2019; Han *et al*. 2021b). Together, these observations indicate that cells in culture often evolve complex genome architectures that deviate substantially from their original source material. Resolving the evolutionary processes that govern the transition from wild-type to complex cell line genome architectures is important for understanding the stability of cell line genotypes and the reproducibility of cell-line-based research. However, the complexity of cell line genomes can impose limitations on efforts to perform *de novo* whole-genome assembly (WGA) (Miller *et al*. 2018a,b; Nattestad *et al*. 2018) and thus limit the ability to study cell line genome structure and evolution using traditional WGA-based bioinformatics approaches.

Like many animal cell lines, Schneider-2 (S2) cells from the model insect *Drosophila* have undergone polyploidization (Schneider 1972; Lee *et al*. 2014), and display substantial small- and large-scale segmental aneuploidy (Zhang *et al*. 2010; Lee *et al*. 2014; Han *et al*. 2021b). In addition, S2 and other *Drosophila* cell lines exhibit a higher abundance of transposable element (TE) sequences compared to whole flies (Potter *et al*. 1979; Ilyin *et al*. 1980; Rahman *et al*. 2015), with TE families that are abundant in S2 cells differing from those amplified in other *Drosophila* cell lines (Echalier 1997; Rahman *et al*. 2015; Han *et al*. 2021a; Mariyappa *et al*. 2021). However, little is known about TE sequence variation in S2 cells or other *Drosophila* cell lines. For example, it is generally unknown whether the proliferation of particular TE families in *Drosophila* cell lines is caused by one or more source lineages (Maisonhaute *et al*. 2007). The lack of understanding about TE sequences in *Drosophila* cell lines is mainly due to previous studies using short-read sequencing data (Rahman *et al*. 2015; Han *et al*. 2021a,b), which typically does not allow complete assembly of TE insertions or other structural variants (Alkan *et al*. 2011; Tattini *et al*. 2015; Kosugi *et al*. 2019; Zhao *et al*. 2021).

Recent advances in long-read DNA sequencing technologies have substantially improved the quality of WGAs, including a better representation of repetitive sequences such as TEs (Berlin *et al*. 2015). In *Drosophila*, long-read WGAs of homozygous diploid genomes such as those from inbred fly stocks can achieve high contiguity and permit detailed analysis of structural variation including TE insertions (Berlin *et al*. 2015; Chakraborty *et al*. 2018; Bracewell *et al*. 2019; Chang *et al*. 2019; Mohamed *et al*. 2020; Ellison and Cao 2020; Hemmer *et al*. 2020; Wierzbicki *et al*. 2021). However, successful WGA using long reads remains limited by complex genome features including polyploidy, heterozygosity, and high repeat content, all of which are present in cell lines such as *Drosophila* S2 cells (Schneider 1972; Potter *et al*. 1979; Ilyin *et al*. 1980; Zhang *et al*. 2010; Lee *et al*. 2014; Rahman *et al*. 2015; Han *et al*. 2021a). In fact, the state-of-the-art long-read assemblies of wild-type diploid genomes still suffer from the presence of repeats and heterozygosity, which may result in assembly gaps and haplotype duplication artifacts (Rhie *et al*. 2021; Peona *et al*. 2021). Therefore, assembly of a complex *Drosophila* cell line genome is likely to result in substantially more fragmented WGAs than those generated from homozygous diploid fly stocks (Fig. 1), and this degradation of assembly quality could impact the subsequent analysis of TE sequences.

**Figure 1.**
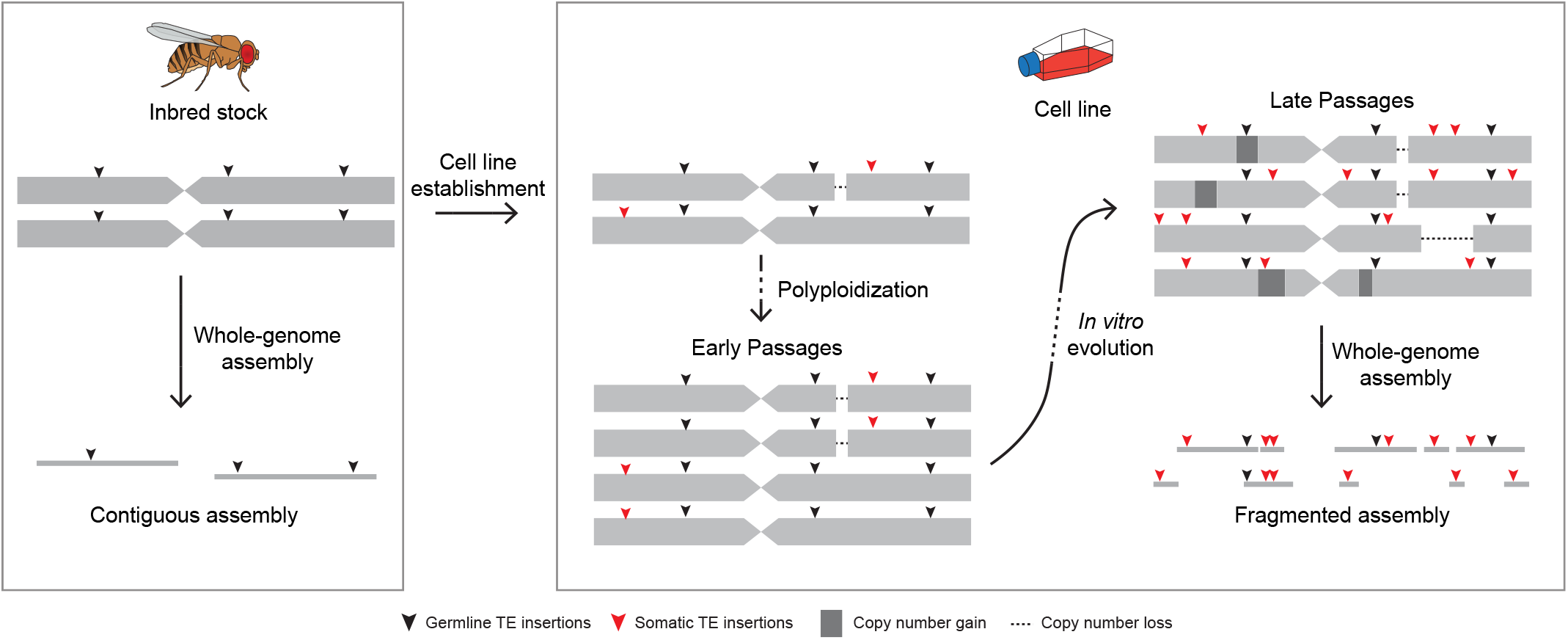
Genome architecture complexity hinders whole-genome assembly in long-term cultured cell lines. The inbred fly stock has diploid genome that includes homozygous variations, which allows contiguous whole-genome assembly (WGA). In comparison, cell lines established from inbred fly stock undergo polyploidization and accumulates heterozygous variations including segmental aneuploidy and haplotype-specific TE insertions during long-term culture. The complexity of polyploid genome with heterozygous variants may lead to highly fragmented WGA and as a result limit the utility of using WGA to study TE sequence evolution.

To gain better insight into the role of transposition during genome evolution in animal cell culture, here we sequenced the genome of a commonly-used variant of S2 cells, the S2R+ cell line (Yanagawa *et al*. 1998), using PacBio long-read and 10x Genomics linked-read technologies. As predicted, WGAs of S2R+ from long-read sequencing data were highly fragmented and yielded highly variable estimates of TE content using different assembly methods. To circumvent the limitations of WGA and characterize TE content in *Drosophila* cell lines, we developed a novel TE detection tool called TELR (<u>T</u>ransposable <u>E</u>lements from <u>L</u>ong <u>R</u>eads, pronounced “Teller”) that can predict non-reference TE insertions based on a long-read sequence dataset, reference genome, and TE library. Importantly, TELR can detect haplotype-specific TE insertions, reconstruct TE sequences, and estimate intra-sample TE allele frequencies (TAFs) from complex genomes that are not amenable to WGA. We applied TELR to our PacBio long-read dataset for S2R+ and similar datasets for a geographically-diverse panel of *D. melanogaster* inbred fly strains from the *Drosophila* Synthetic Population Resource (DSPR) (Chakraborty *et al*. 2019). We discovered a large number of haplotype-specific TE insertions from a subset of LTR retrotransposon families in the tetraploid S2R+ cell line. We inferred that these haplotype-specific insertions came from somatic transposition events that occurred *in vitro* after initial cell line establishment and subsequent tetraploidization (Schneider 1972; Lee *et al*. 2014). We also performed phylogenomic analysis on the full-length TE sequences that were assembled by TELR, which revealed that amplification of TE families in *Drosophila* cell lines can be caused by activity of one or multiple source lineages. Together, our work provides a novel computational framework to study polymorphic TEs in complex heterozygous or polyploid genomes and improves our understanding of the mechanisms of genome evolution during long-term animal cell culture.

## Results

### Fragmented assemblies yield variable estimates of TE content in the S2R+ genome

To better understand the process of TE amplification in the S2R+ cell line genome, we initially sought to use a *de novo* assembly-based approach by generating PacBio long-read (132X average depth) and 10x Genomics linked-read (89X average depth) sequencing data and assembled these data using a variety of state-of-the-art WGA software (Bankevich *et al*. 2012; Chin *et al*. 2016; Koren *et al*. 2017; Weisenfeld *et al*. 2017; Ruan and Li 2020; Kolmogorov *et al*. 2019). All S2R+ whole-genome assemblies (WGAs) using long reads (Canu, FALCON-Unzip, wtdbg2, and Flye) or linked reads (Supernova) had better contiguities compared to a SPAdes assembly of standard Illumina paired-end short read data (Fig. 2A; Table S1). However, S2R+ WGAs from different sequencing technologies and assemblers varied substantially in their contiguities and levels of duplicated BUSCOs (Fig. 2A,B; Table S1). Canu assembly of the S2R+ PacBio data displayed the highest level of BUSCO duplication and the longest total assembly length (Fig. 2B; Table S1). We speculated that the high degree of BUSCO duplication in the Canu S2R+ assembly could be caused by haplotype-induced duplication artifacts in a partially-phased assembly that contained contigs from multiple haplotypes of the same locus (Kelley and Salzberg 2010; Dias *et al*. 2021). To test this, we took advantage of the fact that FALCON-Unzip leverages structural variants to phase heterozygous regions into a primary assembly (“FALCON-Unzip_p”) and alternative haplotigs (Chin *et al*. 2016). Similar to the Canu assembly, combining the primary FALCON-Unzip assembly with alternative haplotigs (“FALCON-Unzip_ph”) resulted a higher level of BUSCO duplication (Fig. 2B). This result suggested that many regions of the S2R+ genome contain haplotype-specific structural variants that can lead to secondary haplotigs (and haplotype-induced BUSCO duplication) in the Canu and Falcon-Unzip assemblies.

**Figure 2.**
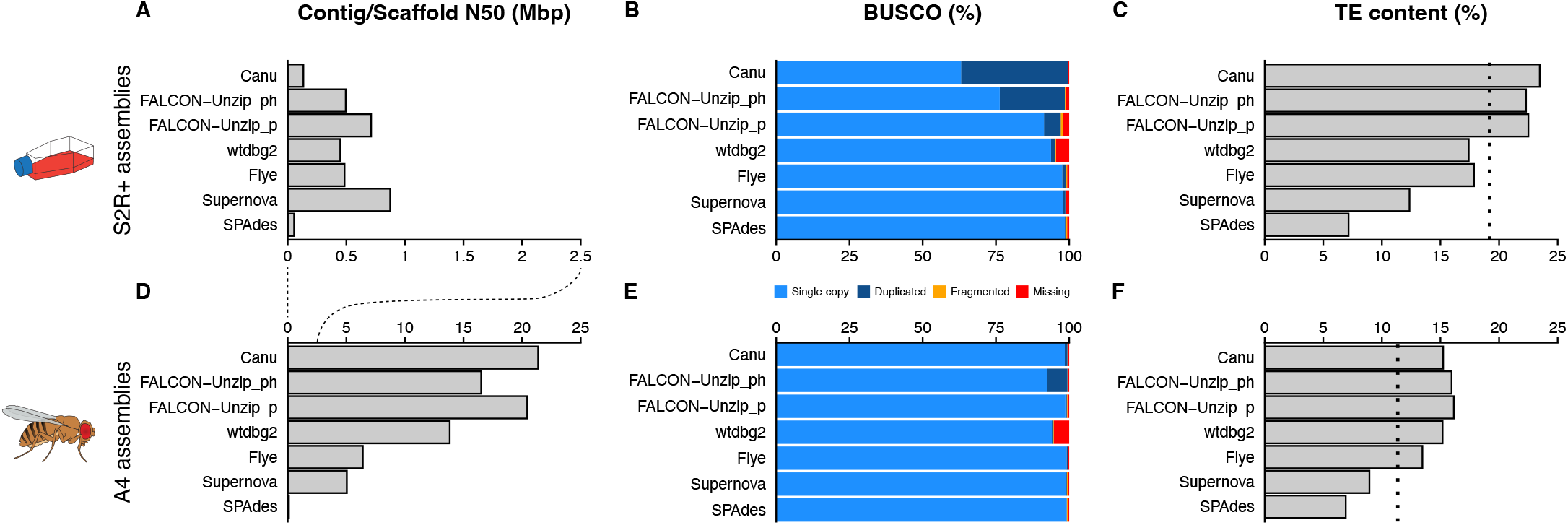
Lower contiguity, and higher BUSCO duplication and TE content in whole-genome assemblies of S2R+ compared to those from an inbred fly strain. (A) and (D) include contig (Canu, FALCON-Unzip, and wtdbg2) and scaffold (Flye, Supernova, and SPAdes) N50 values for S2R+ and A4 whole-genome assemblies, respectively. (B) and (E) include BUSCO (Benchmarking Universal Single-Copy Orthologs) analysis with the Diptera gene set from OrthoDBv10 on S2R+ and A4 assemblies, respectively. (C) and (F) include RepeatMasker estimates of TE content in WGAs of S2R+ and A4, respectively. Dotted lines in (C) and (F) represent RepeatMasker estimates of TE content from raw Illumina reads. “FALCON-Unzip_p” represents primary contigs, “FALCON-Unzip_ph” represents primary contigs + haplotigs. Note that the scale bar is different in (A) and (D).

N50s for all S2R+ WGAs were less than 1 Mbp, which is more than ten-fold smaller than the size of assembled chromosome arms in the *Drosophila* reference genome (Hoskins *et al*. 2015). To assess how S2R+ cell line WGAs compared to those from whole flies of inbred stocks, we also generated WGAs for a highly inbred *D. melanogaster* strain called A4 using available PacBio long-read data (110x average depth) from Chakraborty *et al*. (2019) and a 10x Genomics linked-read dataset for A4 generated in this study (118X average depth) using identical assembly software and parameters as for S2R+. We found that WGAs for A4 have reference-grade contiguities and exhibit lower variation in levels of BUSCO duplication than WGAs for the S2R+ cell line (Fig. 2D,E; Table S2). Given that the A4 strain is diploid homozygous (Chakraborty *et al*. 2019), these results suggest that the highly fragmented WGAs for S2R+ are likely caused by polyploidy, aneuploidy, or heterozygosity in the S2R+ cell line genome rather than limitations of current sequencing or assembly methods.

In addition to assembly quality, estimates of TE content in WGAs varied substantially for both S2R+ and A4 (Fig. 2C,F; Table S1 and S2). Compared to unbiased estimates of TE content based on RepeatMasker analysis of unassembled short reads (dotted lines in Fig. 2C,F) (Sackton *et al*. 2009), long-read WGAs for both the S2R+ and A4 genomes typically gave similar or higher estimates of TE content, while short read WGAs always gave lower estimates. In particular, the Canu and Falcon-Unzip assemblies that we infer include alternative haplotigs gave the highest estimates of TE content relative to unassembled short read data, suggesting the possibility of haplotype-specific TE insertions in these assemblies. In addition to differences in overall TE content, we observed higher variation in the abundance of different TE families across sequencing and assembly technologies in WGAs for S2R+ (Fig. S1A) compared to A4 (Fig. S1B), indicating that WGA-based inferences about TE family abundance in S2R+ are highly dependent on sequencing and assembly technology. Despite this variation, higher estimates of overall TE content were observed in S2R+ WGAs relative to A4 WGAs for all sequencing or assembly technologies used (Fig. 2C,F; Table S1 and S2). However, because of the relatively poor quality and high variation in estimates of TE content among WGAs generated from S2R+ long-read and linked-read data, we concluded that an alternative WGA-independent approach that is better suited to the complexities of cell line genome architecture was necessary to reliably study TE content in S2R+ cells.

### A novel long-read bioinformatics method reveals TE families enriched in S2R+ relative to wild type Drosophila strains

To circumvent the impact of fragmented WGAs on the analysis of TE content in complex cell line genomes, we developed a new TE detection method called “TELR” (<u>T</u>ransposable <u>E</u>lements from <u>L</u>ong <u>R</u>eads; https://github.com/bergmanlab/telr) that allows the identification, assembly, and allele frequency estimation of non-reference TE insertions using long-read data (Fig. 3). Briefly, TELR first aligns long reads to a reference genome to identify insertion variants using Sniffles (Sedlazeck *et al*. 2018). The general pool of insertion variants identified by Sniffles is then filtered by aligning putative insertion sequences to library of curated TE sequences to identify candidate TE insertion loci. For each candidate TE insertion locus, TELR then performs a local assembly using all reads that support the putative TE insertion event. Finally, TELR annotates TE sequence in each assembled contig, predicts the precise location of the TE insertion on reference co-ordinates, then remaps all reads in the vicinity of each insertion to the assembled TE contig to estimate TAF (see Materials and Methods for details).

**Figure 3.**
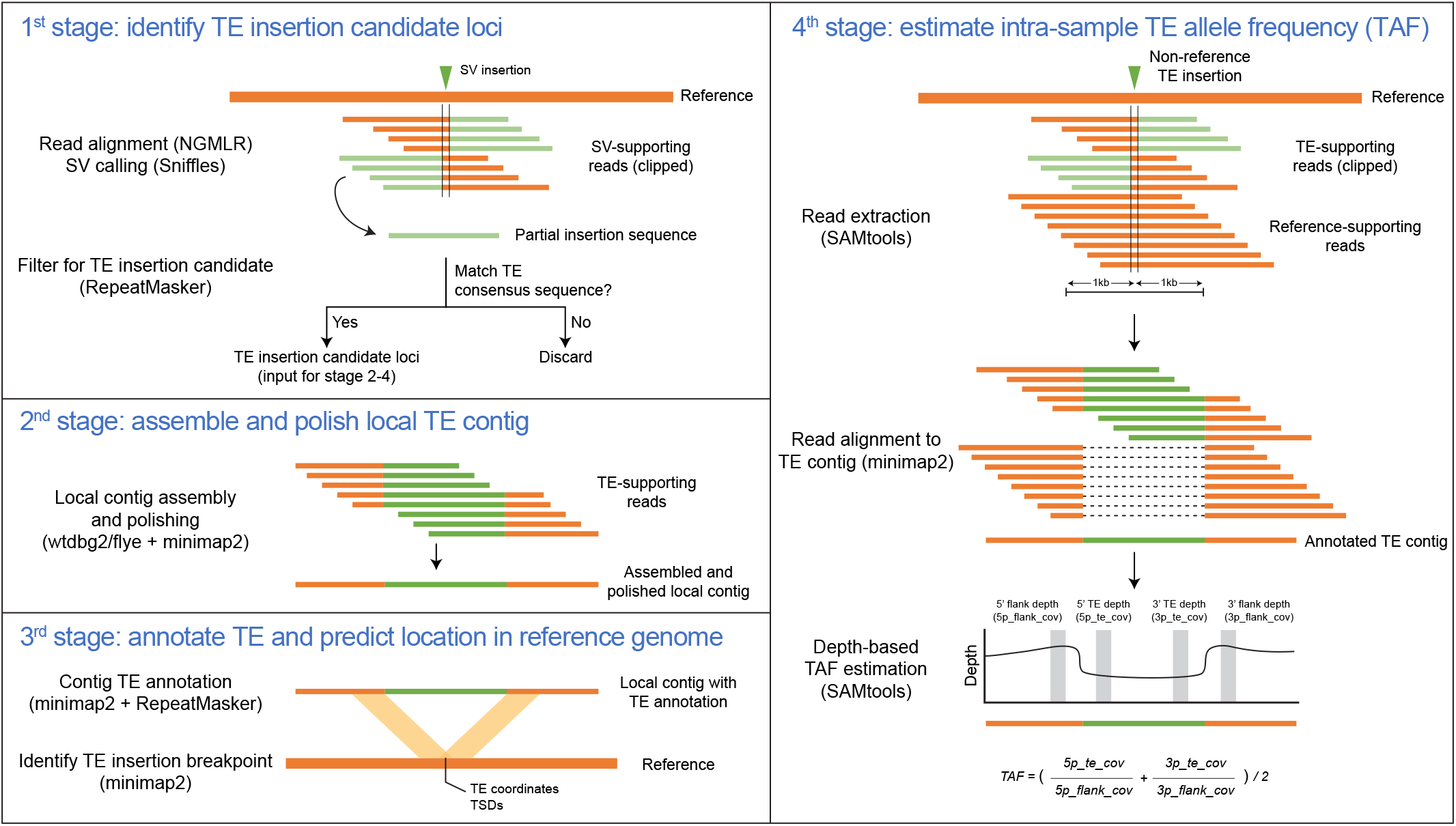
TELR workflow to predict non-reference TE and estimate intra-sample allele frequency. TELR is a non-reference transposable element (TE) detector from long read sequencing data. The TELR pipeline consists of four main stages. In the first stage, TELR aligns long reads to a reference and identify insertions using Sniffles (Sedlazeck *et al*. 2018). TELR then screens for non-reference TE insertion candidate locus by computing nucleotide similarity between partial insertion sequence provided by Sniffles and TE consensus sequences. In the second stage, TELR use SV-supporting reads from Sniffles to assemble and polish local contig using wtdbg2 (Ruan and Li 2020), flye (Kolmogorov *et al*. 2019), and minimap2 (Li 2018). In the third stage, The TE boundaries and family are annotated in the local contig using minimap2 and RepeatMasker, and the TE flanking sequences are used to determine the TE coordinates and target-site duplications by mapping to the reference genome with minimap2. In the fourth stage, TELR determines the intra-sample allele frequency of each TE insertion by extracting all reads in a 2kb span around the insertion locus and aligning them to the TE contig. The mapped read depth over TE and flanking sequences are then used to calculate the intra-sample TE allele frequency (TAF).

Using TELR we identified 2,402 non-reference TE insertions in euchromatic regions of the S2R+ genome, which is a ∼5-fold increase relative to the number identified in A4 (n=490; Fig. 4A). These overall differences in non-reference TE abundance between S2R+ and A4 are unlikely to be caused by variation in coverage and read length between the S2R+ and A4 datasets, as shown by analysis of read length and coverage normalized datasets for S2R+ and A4 (Fig. S2). Despite a drop in the number of predictions in the normalized data relative to the full dataset, TELR still predicted substantially more TEs in S2R+ compared to A4 at all coverage levels (Fig. S2). This analysis also revealed that, unlike A4 which plateaued in the number of non-reference TE insertions at a normalized read depth of 50X, detection of non-reference TEs in S2R+ is likely not saturated even at 75X. Therefore, in order to maximize TE prediction sensitivity, we used the complete non-normalized Pacbio data for S2R+ and all whole-fly strains in subsequent analyses.

**Figure 4.**
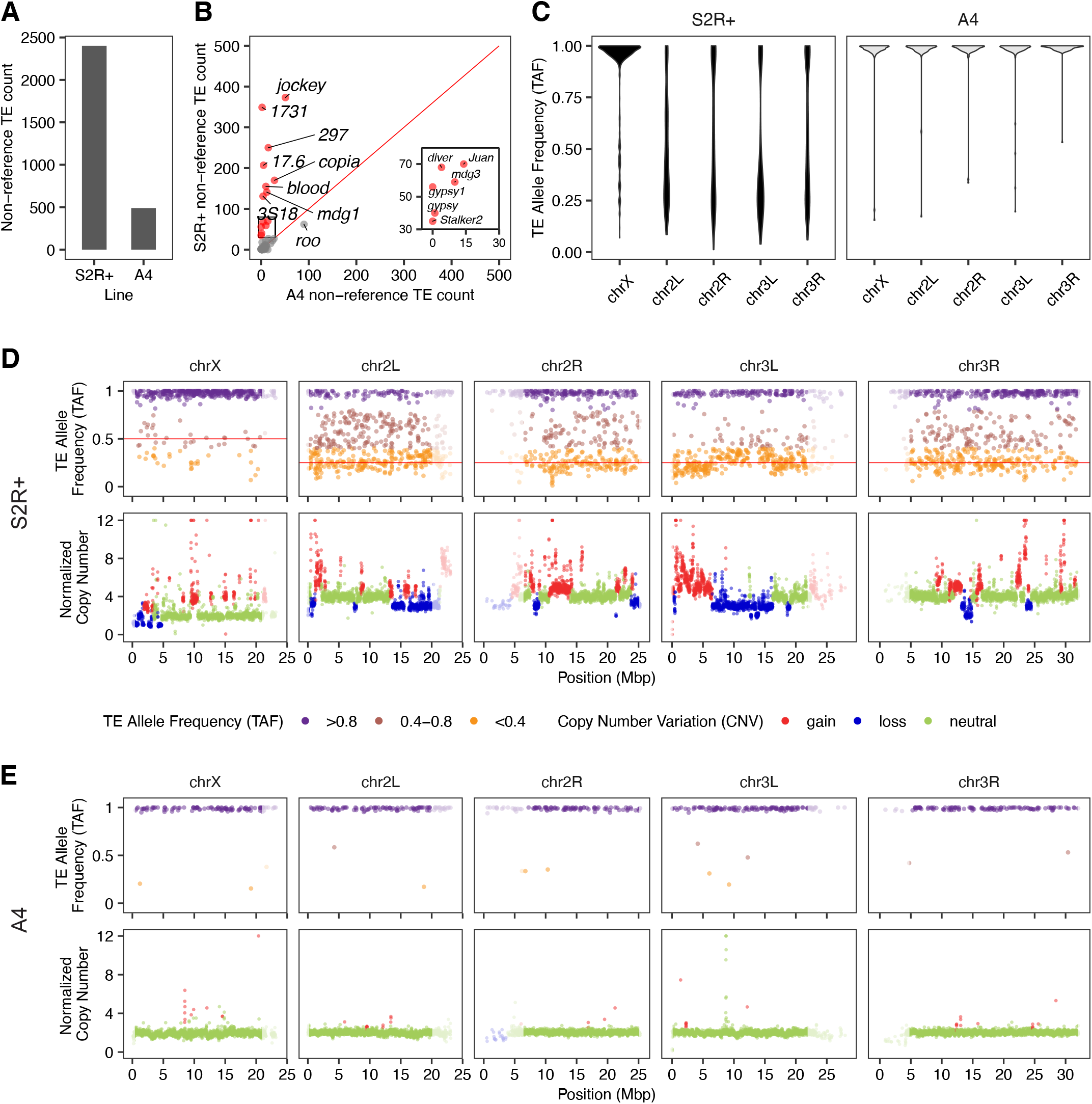
Long-read non-reference TE prediction with TELR reveals multiple families amplified during cell culture. **A** Total number of non-reference TE predictions made by TELR for S2R+ and A4. **B** Number of non-reference TE predictions made by TELR for S2R+ and A4 separated by families with the 14 most abundant families in S2R+ highlighted in red. The insert box is a zoomed plot that includes 6 abundant families in S2R+. **C** TAF distribution by chromosome arm for S2R+ and A4. **D-E** Genome-wide TAF and copy number profiles for S2R+ (D) and A4 (E). Low recombination regions are shaded in grey.

Partitioning the number of non-reference TE insertions predicted by TELR in the complete S2R+ and A4 PacBio datasets by TE family revealed a subset of 14 TE families that are enriched in S2R+ relative to A4 (Fig. 4B; Fig. S5). These S2R+-specific TE families consist mostly of long terminal repeat (LTR) retrotransposons with the exception of *jockey* and *Juan*, which are non-LTR retrotransposons (Fig. 4B; Fig. S5). The TE families revealed by TELR to be enriched in S2R+ relative to A4 were independently cross-validated using short-read sequences and two independent short-read TE detection methods (Fig. S3) (Han *et al*. 2021a; Zhuang *et al*. 2014).

We next used TELR to predict non-reference TEs in PacBio datasets for 13 geographically-diverse *D. melanogaster* inbred strains (including A4) from the DSPR project (Chakraborty *et al*. 2019). This analysis revealed that S2R+ has more non-reference TE insertions than any of the DSPR strains surveyed (range: 445-658; Fig. S4). Partitioning TELR predictions by TE family reveals that only eight TE families account for ∼75% of non-reference insertions in S2R+, most of which are LTR retrotransposons (Fig. S4; Fig. S5). In comparison, 10-16 TE families contribute ∼75% of all non-reference TE insertions in each of the DSPR strain, and they represent a more balanced distribution of LTR retrotransposons, non-LTR retrotransposons, and DNA transposons (Fig. S4; Fig. S5). We also observed strain-specific TE expansions, which we define as a greater than 3-fold increase in the number of non-reference TE insertions for a specific family relative to the mean values across all strains. For example, we see strain-specific expansions of *1360* (n=23, mean=7.13) in A2 (from Colombia), *hopper* (n=114, mean=18.4) in A6 (from USA), as well as *Doc* (n=113, mean=26.5) and *Quasimodo* (n=28, mean=7) in B2 (from South Africa) (Fig. S5).

### Accurate estimation of intra-sample allele frequencies supports haplotype-specific TE insertion after tetraploidy in the S2R+ genome

An important feature of the TELR system is the ability to estimate the intra-sample allele frequency of non-reference TE insertions (Fig. 3), which allowed us to observe drastic differences between S2R+ and A4 in genome-wide TAF patterns. TE insertions in S2R+ display a wide range of allele frequencies, with a striking difference in TAF distributions on the X chromosome relative to the autosomal arms (Fig. 4C; Fig. 4D). In contrast, non-reference TEs in the highly-inbred strain A4 (King *et al*. 2012) are mostly enriched at TAF values ∼1 on all chromosome arms (Fig. 4C; Fig. 4E). Broad-scale patterns of TAF distributions across the S2R+ and A4 genomes detected by TELR using long-read sequences were independently cross-validated using short-read sequences and two independent short-read TE detection methods (Fig. S6) (Han *et al*. 2021a; Zhuang *et al*. 2014).

Like A4, non-reference TEs in other DSPR strains are mostly homozygous with TAF values enriched at the expected value of ∼1 for highly inbred diploid fly stocks (Fig. S7). However, our TELR analysis of DSPR datasets revealed two striking exceptions to this pattern. First, A2 displays mostly heterozygous TE insertions across chromosome arm 3R, which coincides with the presence of a known heterozygous chromosomal inversion in this strain (*In(3R)P*) that prevents full inbreeding (King *et al*. 2012). Second, TAF values in A7 are enriched at ∼0.25 and ∼0.75 across the whole genome (Fig. S7). This TAF pattern is unusual since A7 is thought to be fully inbred and devoid of large chromosomal inversions (King *et al*. 2012). We hypothesized that the bimodal TAF profile in A7 could be indicative of contamination in the A7 data with PacBio reads from a different fly strain in the DSPR project. Indeed, intersecting TELR predictions between A7 and other DSPR strains revealed an unusually large number of non-referenece TE insertion overlaps between strains A7 and B3 (Table S3). Moreover, shared TE insertions between A7 and B3 have TAF enriched at ∼0.25, which could be explained by ∼25% of the A7 dataset being contaminated with reads from B3 (Fig S8). Our inference of contamination in the A7 dataset with reads from another DSPR strain can also explain the observations that A7 has the highest number of non-reference TEs in our TELR analysis (Fig S4), and that the A7 WGA reported in Chakraborty *et al*. (2019) has the highest level of BUSCO duplication, longest assembly length, and most scaffolds of all DSPR strains in that study.

In S2R+, we observed a clear enrichment for TE insertions on the autosomes to have TAFs ∼0.25 (Fig. 4C; Fig. 4D), which can be explained by haplotype-specific TE insertions that occurred after initial cell line establishment and subsequent tetraploidization (Fig. 5A) (Schneider 1972; Lee *et al*. 2014). In contrast to the autosomes, TE insertions on the X chromosome in S2R+ are enriched at TAFs ∼1 (Fig. 4C; Fig. 4D). The X chromosome in the tetraploid S2R+ genome has a baseline ploidy of two since the S2 lineage is thought to have been derived from a hemizygous male genotype (Lee *et al*. 2014). Thus, the enrichment of X-chromosome TE insertions with TAF ∼1 could be explained by a recent loss of heterozygosity (LOH) event in the X chromosome of S2R+ through mitotic recombination. This explanation is plausible since a previous study has shown that copy-neutral LOH events in cell culture can shape TAF profiles over large genomic regions in *Drosophila* cell lines (Han *et al*. 2021a).

**Figure 5.**
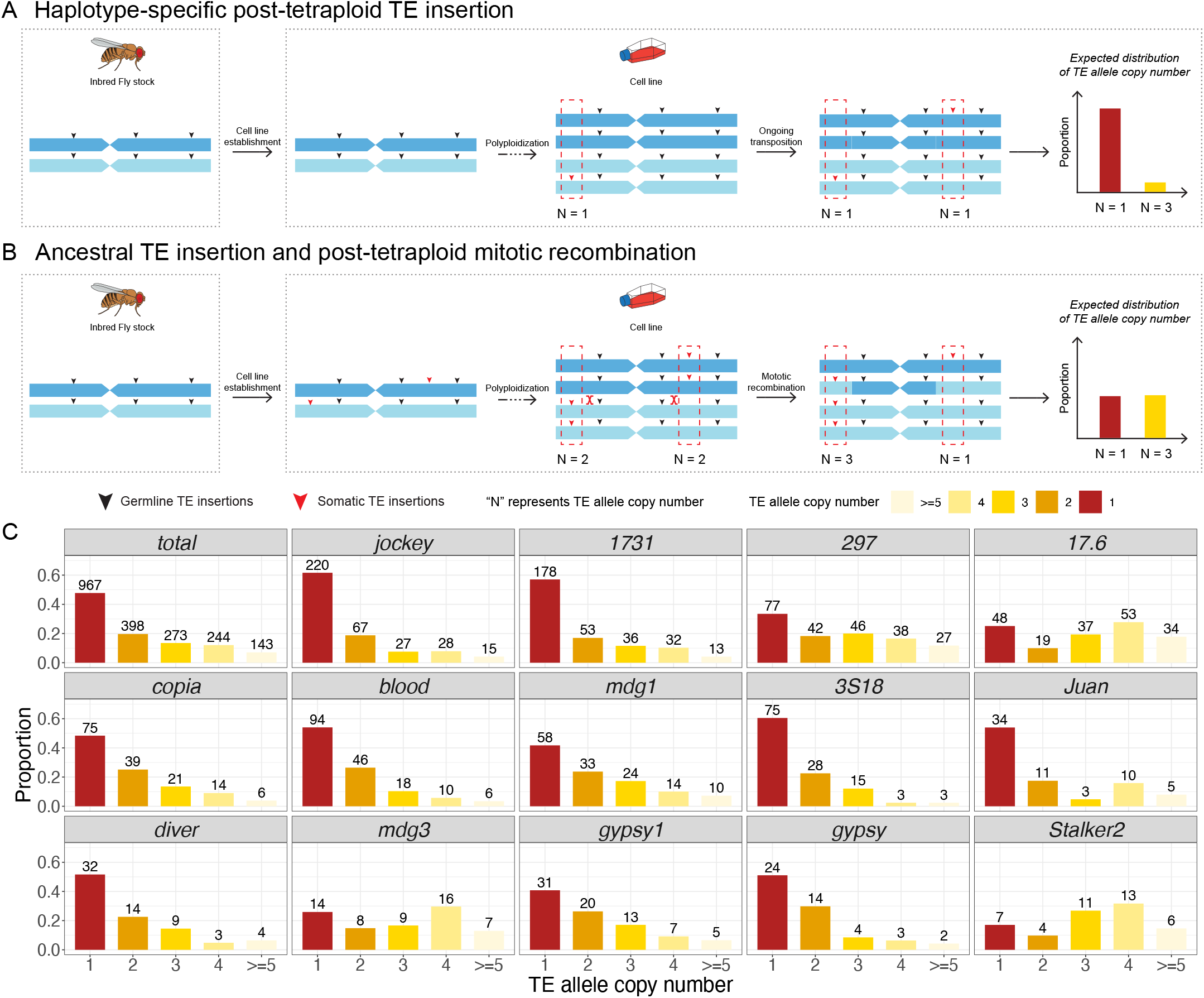
TE allele copy number distribution supports haplotype-specific TE insertion after tetraploidy in the S2R+ genome. **A-B** Two hypotheses that could explain the observation of haplotype-specific TE insertions in the tetraploid S2R+ genome. **C** Distribution on proportion of TE allele copy number for all TEs combined and for 14 TE families that are amplified in S2R+. The TE allele copy number is estimated based on TAF predicted by TELR and local copy number predicted by Control-FREEC (Boeva *et al*. 2012). The histogram is colorized based on TE allele copy number. The number above each bar represents number of TEs under each TE allele copy number category.

Assuming uniform copy number throughout the genome, haplotype-specific autosomal TE insertions that occured in the S2R+ after tetraploidy are expected to have TAFs at ∼0.25. However, the extensive copy number variation observed in the S2R+ genome increases or decreases TAF estimates in affected segments relative to this expected value (Fig. 4D). Additionally, we observed many TE insertions on the S2R+ autosomes that have intermediate TAFs between 0.25 and 1.0, suggesting the possibility of other mechanisms besides haplotype-specific post-tetraploid TE insertion to explain the observed TAF distribution. For example, ancestrally-heterozygous diploid TE insertions (either germline insertions in the Oregon-R lab strain that S2R+ was established from, or somatic insertions in the pre-tetraploid stage of S2) could have undergone mitotic recombination events in the post-tetraploid state changing one haplotype from TE-present to TE-absent (Fig. 5B) (Han *et al*. 2021a). Assuming that ancestral heterozygous diploid TE insertions would be randomly distributed on the two different haplotypes of the Oregon-R/pre-tetraploid state of S2R+, these alternative models can be differentiated since mitotic recombination in the post-tetraploid state would have the same probability of increasing or decreasing TE allele copy number (Fig. 5B), whereas haplotype-specific TE insertion would lead to an excess of alleles with a copy number of one (Fig. 5A).

To facilitate the interpretation of TAF values under varying copy number status and more rigorously test the “haplotype-specific post-tetraploid TE insertion” (Fig. 5A) vs “ancestral TE insertion and post-tetraploid mitotic recombination” (Fig. 5B) models, we developed a strategy to predict absolute TE allele copy number for non-reference TE on the autosomes. For each non-reference TE insertion, we multiplied TAF estimates generated by TELR by the local copy number estimated by Control-FREEC (Boeva *et al*. 2012) in regions flanking the TE insertion, then rounded to the nearest integer value. This procedure generated accurate predictions of TE allele copy number on synthetic tetraploid genomes (see Supplemental Text; Fig S9). Our analysis revealed that a significant proportion of non-reference TE insertions from the 14 TE families that are amplified in S2R+ have a predicted TE allele copy number of one (Fig. 5C). Furthermore, we found that number of TEs with predicted TE allele copy number of one is significantly higher than the number of TEs with predicted TE allele copy number of three in autosomal regions of S2R+ overall (Fig. 5C; chi-squared = 388.42, df = 1, p-value < 2.2e-16) and for all but three S2R+-amplified TE families (*mdg3, Stalker2, 17*.*6*). Thus, we conclude that the majority of insertions in TE families that are amplified in S2R+ are caused by haplotype-specific TE insertions that occurred after tetraploidization, rather than ancestral heterozygous insertions that were reduced in copy number after tetraploidization by mitotic recombination.

### TE expansions in Drosophila cell culture can be caused by one or more source lineages

Haplotype-specific TE insertions that occurred after tetraploidization must have occurred somatically during cell culture, and thus provide a rich set of TE sequences to study how TE expansion events occur during *in vitro* genome evolution. For example, it is generally unknown how many source copies or lineages contribute to proliferation of a TE family during cell culture. Using a PCR-based strategy, Maisonhaute *et al*. (2007) previously concluded that all non-reference insertions for the *1731* family in the S2 cell line were derived from a single, strongly-activated source copy. However, only a single TE family was surveyed and the number of *1731* new insertions identified was likely underestimated due to the limitations of the PCR-based strategy in this study. Moreover, it is difficult to conclude whether amplification is due to a single source copy or multiple closely-related copies from a single source lineage. To comprehensively test whether one or more source lineage is responsible for the amplification all 14 TE families that expanded in S2R+ (Fig. 4B), we took advantage of TELR’s ability to assemble non-reference TE sequences and constructed phylogenies using data from S2R+ and 13 whole-fly strains from the DSPR panel (Fig. 6; Fig. S10). Evaluation of TE sequences reconstructed by TELR using simulated datasets suggested that TELR produced high-quality local assemblies (see Supplemental Text; Fig. S11; Fig. S12), and thus can be reliably used to infer the sequence evolution of TEs amplified in the polyploid cell line genomes like S2R+.

**Figure 6.**
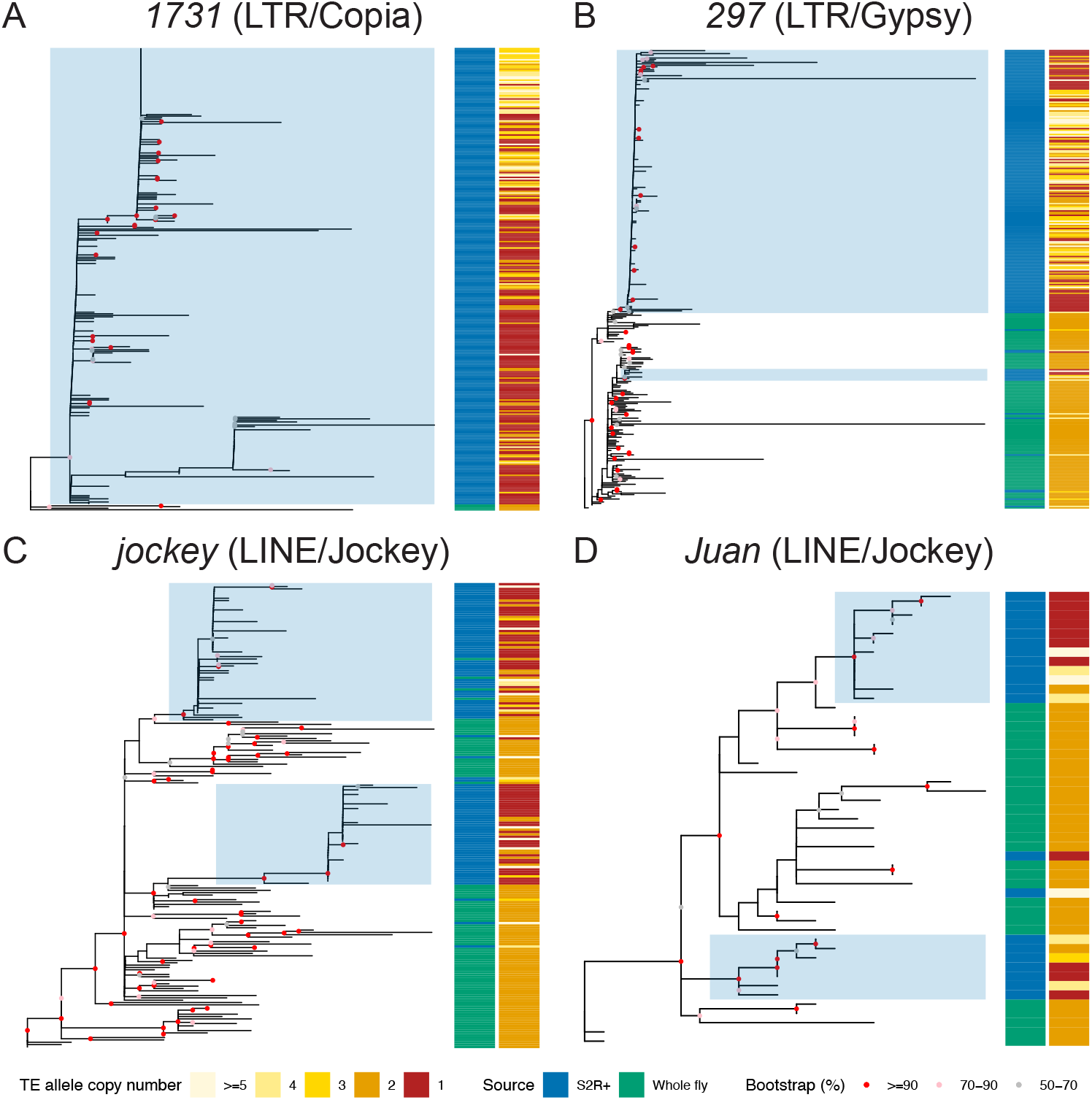
Single and multiple TE source lineage activation in S2R+ cell line. **A-D** Non-reference TE insertion sequences from S2R+ and 11 inbred *Drosophila* fly strains were predicted and assembled by TELR. Only high-quality full-length TE sequences in normal recombination autosomal regions were retained for this analysis (see Materials and Methods for details). TE sequences for each family were aligned using MAFFT (v7.487) (Katoh and Standley 2013). The multiple sequence alignments were used as input in IQ-TREE (v2.1.4-beta) (Minh *et al*. 2020) to build unrooted trees for *1731* (A), *297* (B), *jockey* (C) and *Juan* (D) elements using maximum likelihood approach. The sample source and TE allele copy number were annotated in the sidebars. Blue shading indicates TE expansion event in S2R+ from a single source lineage based on the following criteria: 1) All sequences should form a monophyletic clade, 2) The monophyletic clade should include at least three post-tetraploid cell-line-specific TE insertions, 3) The bootstrap support for the clade should be equal to or higher than 50%, and 4) The proportion of post-tetraploid cell-line-specific TE insertions (i.e. TE allele copy number equal to one) within the clade should be equal to or higher than 20%.

Using the sequences of full-length TE insertions identified by TELR, we designed a set of criteria to identify TE expansion events in S2R+ that start from a single source lineage. First, the TE expansion event should be marked by a monophyletic clade in which ≥30% of TEs are enriched with post-tetraploid insertions in S2R+. Second, the candidate TE expansion clade should have at least 70% bootstrap support. Using these criteria, we annotated TE expansion events in the sequence phylogeny for each of the 14 TE families that are enriched in S2R+ relative to A4 (Fig. 4B, TE families marked in red dots). We only used TE sequences in autosomes for this analysis, given that TE allele copy number distribution in Chromosome X is different from the autosomes presumably due to an LOH event after tetraploidy (see above). We identified a single TE expansion clade for TE families such as *1731, gypsy1, diver, gypsy, mdg3*, and *Stalker2* (Fig. 6; Fig. S10), suggesting that the TE expansion events in the S2R+ cell line for these families came from a single source lineage. We also identified multiple TE expansion clades for TE families such as *jockey, Juan, copia, 3S18*, and *mdg1* (Fig. 6; Fig. S10), suggesting multiple source lineages contribute to the amplification of these families in S2R+. Together, our results revealed that TE expansions in S2R+ can be caused by single or multiple source lineages, and that the pattern of source lineage activation in somatic cell culture is TE family-dependent (Fig. 6; Fig. S10).

## Discussion

Here we report new long-read and linked-read sequence data and develop a novel bioinformatics tool to study the role of transposition during long-term *in vitro* evolution of an animal cell line. Our finding that the complexities of *Drosophila* S2R+ genome architecture preclude the ability to accurately study TE content using long-read or linked-read WGAs motivated the development of a WGA-independent TE detection system called TELR, which can identify, locally assemble, and estimate allele frequency of TEs from long-read sequence data. Our work provides new tools and approaches to study TE biology in complex heterozygous or polyploid genomes found in many other animal cell lines (Lee *et al*. 2014; Nattestad *et al*. 2018; Talsania *et al*. 2019) as well as natural fungal and plant genomes (Todd *et al*. 2017; Meyers and Levin 2006).

Several related WGA-independent bioinformatic methods have recently been developed to detect non-reference TEs using long reads (Disdero and Filee 2017; Jiang *et al*. 2019; Zhou *et al*. 2020; Ewing *et al*. 2020; Chu *et al*. 2021; Kirov *et al*. 2021). These methods use a variety of strategies for TE detection and generate different information for predicted non-reference TEs. Importantly, none of these previously-reported methods for TE detection using long reads can estimate intra-sample TAF, a feature that we implemented in TELR specifically to identify haplotype-specific TE insertions and which enabled our analysis of post-tetraploidy somatic transposition in S2R+. Furthermore, TELR is the only WGA-independent long-read detection tool that outputs a polished assembly of the TE locus, providing a high-quality sequence of both the TE its flanking regions. The polishing step in TELR is especially important to improve sequence quality when using long-read assemblers such as wtdbg2 (Ruan and Li 2020) that do not error correct reads prior to the assembly step. High-quality sequences of predicted TE insertions generated by TELR allowed us to gain the first general insight into the sequence variation underlying TEs proliferation in an animal cell line.

Using the TELR system, we found a significantly higher number of non-reference TEs in S2R+, a sub-line of *Drosophila* S2 cell line, compared to whole fly of highly inbred strain from the DSPR project. The increased TE allele copy number in S2R+ relative to wild type flies is mainly contributed by a subset of mainly LTR and a few non-LTR retrotransposon families. Notably, TE families identified as enriched in S2R+ by TELR using long-read sequences were also detected as having high activity at some point during the history of S2 cell line evolution in an independent analysis of short-read sequences for multiple sublines of S2 cells by Han *et al*. (2021b), providing cross-validation for both approaches. In addition, TELR predicted that a significant proportion of the non-reference TE insertions identified in S2R+ have TE allele copy number of one, which we interpreted as haplotype-specific somatic insertions that occurred after S2R+ cells became tetraploid, subsequent to the initial establishment of the original S2 cell line (Schneider 1972). This interpretation is consistent the main conclusion from Han *et al*. (2021b) that TE amplification in *Drosophila* S2 cells is an ongoing, episodic process rather than being driven solely by an initial burst of transposition dureing cell line establishment. Finally, the phylogenomic analysis using TELR-assembled sequences for TE families enriched in S2R+ suggested that the TE expansion in cell culture could come from a single or multiple source lineages, providing the first general insight into the sequence evolution of TE family expansions in animal cell culture.

## Materials and Methods

### Cell culture

An initial sample of S2R+ cells, which we define as passage 0, was obtained from a routine freeze of cells made by the *Drosophila* RNAi Screening Center (DRSC). Cells from passage 0 were defrosted and recovered in Schneider’s *Drosophila* medium (Thermo) containing 10% FBS (Thermo) and 1X Penicillin-Streptomycin (Thermo), then expanded continually for two additional passages in T75 flasks. Aliquots of cells from passage 3 flasks were frozen, and the remaining cells were expanded to 10 T75 flasks (passage 4A). Passage 4A cells were pooled and harvested to make DNA for PacBio libraries. A frozen stock was defrosted and expanded for two additional passages (passages 4B-5B). Passage 5B cells were harvested to make DNA for 10x Genomics libraries. The provenance of the cell line samples used in this study is depicted in Fig. S13.

### Fly stocks

A stock of *D. melanogaster* strain A4 from the Drosophila Synthetic Population Resource (DSPR) (King *et al*. 2012) was obtained from Stuart Macdonald (University of Kansas) and reared on Instant Drosophila Medium (Carolina Biological, Cary NC) until DNA extraction.

### PacBio library preparation and sequencing

Cells from ten confluent T75 flasks from passage 4A were scraped into a 15mL Falcon tube and centrifuged at 300 x g for 3 min. The pellet was washed in 10 mL of 1X PBS, then resuspended in 7 mL of 1X PBS containing 35 uL of 10 mg/mL RNAse A (Sigma). 200 uL of resuspended cells were aliquoted to 32 Eppendorf tubes containing 200 uL of buffer AL from the Qiagen Blood & Tissue kit, mixed gently by inversion, and incubated at 37 °C for 30 min. 20 uL of Proteinase K solution from the Qiagen Blood & Tissue kit was then added to each tube and mixed gently by inversion. One volume of phenol:chloroform:isoamyl alcohol (24:24:1) was then added and inverted gently to mix for 1 min. Tubes were then spun for 5 min at 21,000 x g. 180 uL of the upper aqueous phase were then removed from each tube, and pairs of tubes were combined. 400 uL of chloroform was then added to each of the 16 tubes, shaken for 1 min to mix, and spun at max speed for 5 min. The top 300 uL was removed and pairs of tubes were combined. 600 uL of chloroform was added to each of the eight tubes, gently inverted 10 times to mix, and then spun at max speed for 5 min. 400 uL of the aqueous phase was removed and pairs of tubes were combined. 1/10 volume of 3M NaOAc was added to each of the four tubes, the remained of the tube was filled with absolute ethanol and then placed at −20 °C overnight. Tubes were then spun 21,000 x g at 4 °C for 15 min, and the supernatant was decanted over paper towels. 70% ethanol was then added to tubes, the pellet was gently resuspended with a P1000 tip, and then placed on ice for 10 min. Tubes were then spun 21,000 x g at 4 °C for 15 min, and the supernatant was decanted over paper towels. The pellet was then resuspended in 50 uL of Buffer EB from the Qiagen Blood & Tissue kit, and gently pipetted with a P200 tip 5 times to resuspend. Purified S2R+ DNA was then used to generate PacBio SMRTbell libraries using the Procedure & Checklist 20 kb Template Preparation using BluePippin Size Selection protocol. The SMRTbell library was sequenced using 31 SMRT cells on a PacBio RS II instrument with a movie time of 240 minutes per SMRT cell, generating a total of 3,510,012 reads (∼28.5 Gbp).

### 10x Genomics library preparation and sequencing

Genomic DNA extraction followed the 10x “Salting Out Method for DNA Extraction from Cells” protocol (https://support.10xgenomics.com/permalink/5H0Dz33gmQOea02iwQU0iK) adapted from Miller *et al*. (1988). Genomic DNA for *D. melanogaster* strain A4 linked-read library was obtained from a single female fly following the 10x Genomics recommended protocol for DNA purification from single insects (https://support.10xgenomics.com/permalink/7HBJeZucc80CwkMAmA4oQ2). Purified DNA was precipitated by addition of 8 mL of ethanol and resuspended in TE buffer and size was analyzed by TapeStation (Agilent) prior to library preparation. Linked-read libraries were then prepared for both S2R+ and A4 after DNA size selection with BluePippin to remove fragments shorter than 15 kb. Libraries were prepared following the 10x Genomics Chromium Genome Reagent Kit Protocol v2 (RevB) using a total DNA input mass of 0.6 ng for each sample. The linked-read libraries were sequenced on an Illumina NextSeq 500 instrument mid-output flow cell with 150 bp paired-end layout, generating 95,280,430 reads for S2R+ (∼13.3 Gbp) and 127,009,398 reads for A4 (∼17.7 Gbp).

### Whole-genome assembly and QC

Raw PacBio reads from S2R+ (generated here; SRX7661404) and A4 from Chakraborty *et al*. (2018) (SRX4713156) were independently used as input for whole-genome assembly with Canu (v2.1.1; genomeSize=180m corOutCoverage=200 “batOptions=-dg 3 -db 3 -dr 1 -ca 500 -cp 50” -pacbio-raw), FALCON-Unzip (pb-falcon v0.2.6; seed coverage = 30, genome_size = 180000000), wtdbg2 v2.5 (-x rs -g 180m), and Flye (v2.8.2) (Chin *et al*. 2016; Koren *et al*. 2017; Kolmogorov *et al*. 2019; Ruan and Li 2020). The reads were re-aligned to the resulting assemblies with pbmm2 (v1.3.0; --preset SUBREAD --sort) and the assemblies were polished with the Arrow algorithm from GenomicConsensus (v2.3.3) using default parameters. FALCON-Unzip performs read re-alignment and Arrow polishing automatically as part of its phasing pipeline.

10x Genomics linked-reads generated here were used as input for whole-genome assembly with Supernova (v2.1.1) for S2R+ (--maxreads=61508497) and A4 (--maxreads=77907944) (Weisenfeld *et al*. 2017). The optimal --maxreads parameter was calculated by Supernova in a previous run to avoid excessive coverage. Supernova assemblies were exported in pseudohap2 format and pseudo-haplotype1 was analyzed.

10x Genomics reads from S2R+ and A4 were also barcode-trimmed with LongRanger (v2.2.2; basic pipeline) (Zheng *et al*. 2016) to create standard paired-end reads as input to SPAdes (v3.15.0) using default parameters (Bankevich *et al*. 2012).

All assemblies were filtered to remove redundancy using the sequniq program from GenomeTools (v1.6.1) (Gremme *et al*. 2013). General assembly statistics were calculated with the stats.sh utility from BBMap (v38.83) (Bushnell 2014). Assembly completeness was assessed with BUSCO (v4.0.6) (Simao *et al*. 2015; Waterhouse *et al*. 2018) and the Diptera ortholog set from OrthoDB (v10) (Kriventseva *et al*. 2019).

### Assessment of overall TE content

Transposable elements were annotated in all WGAs with RepeatMasker (v4.0.7; -s -no_is -nolow -x -e ncbi) (https://www.repeatmasker.org/RepeatMasker/) using v10.2 of the curated library of *D. melanogaster* canonical TE sequences (https://github.com/bergmanlab/transposons). TE abundance was calculated from RepeatMasker .out.gff files as the percentage of bases masked in each assembly.

Barcode-trimmed linked-reads were also used as an assembly-free estimate of TE content in S2R+ and A4. Reads were filtered for adapters and low quality bases, and trimmed to 100 bp using fastp (v0.20.0; --max_len1 100 --max_len2 100 - -length_required 100) (Chen *et al*. 2018). A random sample of 5 million read pairs (10 million reads) was extracted for each dataset using seqtk (v1.3; -s2) (https://github.com/lh3/seqtk) and masked using RepeatMasker (v4.0.7; -s -no_is -nolow –x -e ncbi) and the *D. melanogaster* canonical TE set (v10.2; https://github.com/bergmanlab/transposons). Abundance for each TE family was calculated as the percentage of read bases that were RepeatMasked.

### Detection of non-reference TE insertions using long reads

The TELR pipeline consists of four main stages: (1) general SV detection and filter for TE insertion candidate, (2) local reassembly and polishing of the TE insertion, (3) identification of TE insertion coordinates, and (4) estimation of intra-sample TE insertion allele frequency.

In stage 1, long reads are aligned to the reference genome using NGMLR (v0.2.7) (Sedlazeck *et al*. 2018). The alignment output in BAM format is provided as input for Sniffles (v1.0.12) to detect structural variations (SVs) (Sedlazeck *et al*. 2018). TELR then filters for TE insertion candidates from SVs reported by Sniffles using following criteria: 1) The type of SV is an insertion, 2) The insertion sequence is available, and 3) The insertion sequences include hits from user provided TE consensus library using RepeatMasker (v4.0.7; http://www.repeatmasker.org/).

In stage 2, reads that support the TE insertion candidate locus based on Sniffles output are used as input for wtdbg2 (v2.5) to assemble local contig that covers the TE insertion for each TE insertion candidate locus (Ruan and Li 2020). The local assemblies are then polished using minimap2 (v2.20) (Li 2018) and wtdbg2 (v2.5) (Ruan and Li 2020).

In stage 3, TE consensus library is aligned to the assembled TE insertion contigs using minimap2 and used to define TE-flank boundaries. TE region in each contig is annotated with family info using RepeatMasker (v4.0.7). Sequences flanking the TE insertion are then re-aligned to the reference genome using minimap2 to determine the precise TE insertion coordinates and target site duplication (TSD).

In stage 4, raw reads aligned to the reference genome are extracted within a 1kb interval on either side of the insertion break-points initially defined by Sniffles. The reads are then aligned to the assembled polished contig to identify reads that support the non-reference TE insertion and reference alleles, respectively, in following steps: 1) Reads are aligned to the forward strand of the contig, 5’ flanking sequence depth (5p_flank_cov) and 5’ TE depth (5p_te_cov) are calculated. 2) Reads are aligned to the reverse complement strand of the contig, 5’ flanking sequence depth (3p_flank_cov) and 5’ TE depth (3p_te_cov) are calculated. 3) The TE allele frequency is estimated as (5p_te_cov/5p_flank_cov + 3p_te_cov/3p_flank_cov)/2.

TELR (v0.2; revision bb90a5) was applied to the S2R+ PacBio dataset and to a panel of 13 *D. melanogaster* strains from the *Drosophila* Synthetic Population Resource (DSPR) (Bioproject ID PRJNA418342) (Chakraborty *et al*. 2019). The mapping reference used was release 6 of the *D. melanogaster* reference genome (chr2L, chr2R, chr3L, chr3R, chr4, chrX, chrY, chrM) (Hoskins *et al*. 2015) and the TE library was v10.2 of the *D. melanogaster* canonical TE sequence library (https://github.com/bergmanlab/transposons/blob/master/releases/D_mel_transposon_sequence_set_v10.2.fa).

We used BEDTools (v2.29.0) (Quinlan and Hall 2010) to investigate the possibility of contamination of sample A7 with another strain by intersecting TE predictions between A7 and all other DSPR strains.

### Cross-validation of TELR results using short-read methods

To cross-validate results obtained by TELR, we employed two short-read TE detection methods implemented in McClintock (v2.0; revision 93369ef) (Nelson *et al*. 2017) that output TAF values, which include ngs_te_mapper2 (Han *et al*. 2021a) and TEMP (Zhuang *et al*. 2014). Linked-read data obtained for S2R+ and A4 was barcode-trimmed with LongRanger (v2.2.2; basic pipeline) (Zheng *et al*. 2016), de-interleaved, and trimmed to 100bp using fastp (v0.20.0; –max_len1 100 --max_len2 100 --length_required 100) (Chen *et al*. 2018). This data was downsampled to ∼50X mean mapped read depth for S2R+ (74,648,362 reads) and A4 (76,045,544 reads) before being used as input in McClintock to generate non-redundant non-reference TE insertion predictions.

### Construction of phylogenetic trees using TE sequences from TELR

TE sequences predicted, assembled, and polished by TELR on S2R+ and DSPR dataset were filtered for high-quality full length TE sequences using the following criteria: 1) Sequences from A2 were excluded due to potential inversion-induced gain of heterozygosity (see Discussion for details). 2) Sequences from A7 were excluded due to potential sample contamination (see Discussion for details). 3) Sequences from chromosome X were excluded due to lower coverage compared to autosomes and loss of heterozygosity (LOH) events. 4) Exclude sequences from low recombination regions using boundaries defined by Cridland *et al*. (2013) lifted over to dm6 coordinates. Normal recombination regions included in our analyses were defined as chrX:405967–20928973, chr2L:200000–20100000, chr2R:6412495– 25112477, chr3L:100000–21906900, chr3R:4774278–31974278. We restricted our analysis to normal recombination regions since low recombination regions have high reference TE content which reduces the ability to predict non-reference TE insertions (Bergman *et al*. 2006; Manee *et al*. 2018). 5) Only full-length TE elements based on canonical sequences were included. We first calculated the ratio between each TELR sequence length and the corresponding canonical sequence length. Next, we filtered TELR sequences for full-length copies using a 0.75-1.05 ratio cutoff for *297* and 0.95-1.05 ratio cutoff for other TE families. 6) Only sequences with both 5’ and 3’ flanks mapped to reference genome were included. 7) Only sequences from TE insertions with TAF estimated by TELR were included.

TELR sequences from each family were aligned with MAFFT (v7.487) (Katoh and Standley 2013). The multiple sequence alignments (MSAs) were filtered by trimAI (v1.4.rev15; parameters: -resoverlap 0.75 -seqoverlap 80) to remove spurious sequences. The filtered MSAs were used as input to IQ-TREE (v2.1.4-beta; parameters: -m GTR+G -B 1000) (Minh *et al*. 2020) to generate maximum likelihood trees.

## Data Availability

PacBio and 10x Genomics whole genome sequences generated in this project are available in the NCBI SRA database under accession PRJNA604454. WGAs of long-read and linked-read sequence data for the S2R+ and A4 genomes are available in the EBI BioStudies database under accession S-BSST752. Datasets of TE insertions in the S2R+ and DSPR genomes predicted by TELR are available as Supplemental File 1. Datasets of TE insertions in the S2R+ and A4 genomes predicted by TEMP and ngs_te_mapper2 are available as Supplemental File 2. Multiple sequence alignments of TE insertion sequences identified by TELR in the S2R+ and DSPR genomes are available as Supplemental File 3. Tree files for phylogenies of TE insertion sequences identified by TELR in the S2R+ and DSPR genomes are available as Supplemental File 4.

## Acknowledgements

We thank Stuart Macdonald (University of Kansas) for providing fly stocks; Christina McHenry and Robert Lyons at the University of Michigan Biomedical Research Core Facilities for assistance with PacBio library preparation and sequencing; Noah Workman, Julia Portocarrero and Magdy Alabady at the University of Georgia Genomics and Bioinformatics Core for assistance with 10x Genomics library preparation and Illumina sequencing; the Georgia Advanced Computing Resource Center for computing time; members of the Bergman Lab for helpful comments throughout the project; and XX for comments on the manuscript. This work was supported by the Human Frontiers of Science (C.M.B.) and Georgia Research Foundation (C.M.B.) and the Howard Hughes Medical Institute (N.P.).

