## Supplemental Tables and Figures for "Local assembly of long reads enables phylogenomics of transposable elements in a polyploid cell line"

### 1 Supplementary Text

#### 1.1 Evaluate TELR on predicting non-reference TE insertion coordinate and family

We generated synthetic datasets using reads simulated from the ISO1 (dm6) and A4 (GCA_003401745.1) (Chakraborty *et al.*, 2018) genome assemblies to evaluate TELR on predicting non-reference TE insertion under different ploidy, zygosity and coverage settings. In principle, a good predictor should be able to accurately predict "non-reference" insertions that are present in genome 1 (e.g., ISO1) but absent from genome 2 (e.g., A4) using reads simulated from genome 1 and 2 mapped to genome 2. Synthetic datasets under different settings were created as follows: 1) We simulated Pacbio reads from ISO1 to model diploid homozygous insertions. 2) We simulated and combined Pacbio reads from both ISO1 and A4 with equal coverages to model diploid heterozygous insertions. 3) We simulated and combined Pacbio reads ISO1 and A4 with 1:3, 2:2, 3:1, and 4:0 ratio to model tetraploid simplex, duplex, triplex and quadruplex insertions, respectively (Table S5). The simulations were conducted using pbsim2 (v2.0.1; P6C4 HMM model) (Ono *et al.*, 2021) under 50X, 100X, 150X, and 200X coverages for all ploidy, zygosity, and coverage settings. The synthetic datasets were used as input to TELR to detect non-reference TE insertions (v0.2; revision bb90a5a; options: –assembler wtdbg2 –polisher flye -p 1). The A4 assembly was used as the reference genome and the Berkeley *Drosophila* Genome Project canonical TE dataset v10.2 was used for these analyses.

As ground truth for evaluating TELR performance, curated TE annotations from the release 6.38 version of *D. melanogaster* ISO1 genome (`http://ftp.flybase.net/releases/FB2021_01/dmel_r6.38/gff/dmel-all-r6.38.gff.gz`) were lifted over to A4 genome assembly. After excluding *INE-1* insertions, TE insertions in low recombination regions, TEs without TAF predictions, and TEs without flank alignment support for both sides, 1163 curated TEs in ISO1 could be lifted over to A4 on the basis of their flanking regions. TELR predictions were considered true positives if the predicted TE insertion coordinates were within a 5bp window of a lifted over ISO1 TE annotation and if the predicted TE family was the same as the lifted over annotation. The final benchmark results for TELR applied to synthetic datasets are summarized in Table S5. The results suggested that TELR has high precision (≥95%) under all ploidy, zygosity and coverage settings. In contrast, TELR's recall was much lower, especially at low effective coverage levels. These results indicate that the non-reference TE insertion predictions made by TELR are highly accurate, however, the method has an appreciable false negative rate especially when the effective coverage is lower than 50X.

#### 1.2 Evaluation of TE allele copy number estimation in diploid and tetraploid genome using long-read data

We estimated the TE allele copy number by multiplying TAF predicted by TELR and local copy number predicted by Control-FREEC (Boeva *et al.*, 2012). The estimated TE allele copy number is then rounded into integer value. To evaluate the TE allele copy number estimation accuracy, we used synthetic datasets generated from ISO1 and A4 genome assemblies under different coverage, ploidy, and zygosity settings (see details in Section 1.1). Only non-*INE-1*, non-nested insertions from normal recombination regions, with flank alignment support for both sides, and for which the TAF could be calculated were included in this analysis. The benchmark results were summarized in Table S6 and Table S7 for diploid and tetraploid genome, respectively. Our estimated TE allele copy number had over 99.4% precision for diploid genomes and over 89.7% precision for tetraploid genomes under all coverage levels.

#### 1.3 Evaluate TELR on TE sequence quality

Ideally, TELR should produce local contig assemblies that can be perfectly aligned to the corresponding TE loci in the ISO1 genome assembly. To evaluate quality of local contigs and TE sequences assembled and polished by TELR, we used synthetic datasets generated from ISO1 and A4 genome assemblies under different coverage, ploidy, and zygosity settings (see details in Section 1.1). Only non-*INE-1*, non-nested insertions from normal recombination regions, with flank alignment support for both sides, and for which the TAF could be calculated were included in this analysis. For a given TELR run, each reported TE sequence plus 500bp flanks upstream and downstream of TE region in the local contig assembly (later referred to as "TELR TE locus") was aligned to the ISO1 genome assembly. All TELR TE loci can be uniquely aligned to ISO1 (Figure S11). Next, for each TE locus predicted by TELR that have a corresponding curated TE annotation, we compared the TELR TE sequence with corresponding TE sequence in ISO1 based on curated TE annotation. The nucleotide identity between TELR TE sequences and corresponding TE sequences in ISO1 are summarized in Figure S12. Note that although all TELR TE loci can be aligned to ISO1, a subset of TELR TE loci don't have corresponding curated TE annotation (mean = 29.1, sd =

5.14 for number of unmatched TELR TE loci across all TELR runs), which can be explained by missing info in the curated TE annotation.

#### 2 Supplementary Tables

Table S1: **Statistics for S2R+ genome assemblies.** FALCON-Unzip_p = primary contigs; FALCON-Unzip_h = haplotigs; FALCON-Unzip_ph = primary + haplotigs.

| | Canu | Falcon-Unzip_p | Falcon-Unzip_h | Falcon-Unzip_ph | wtdbg2 | Flye | Supernova | SPAdes |
|---|---|---|---|---|---|---|---|---|
| Assembly size (bp) | 288,352,585 | 166,182,448 | 43,221,666 | 209,404,114 | 146,747,034 | 144,559,338 | 136,942,585 | 138,619,786 |
| Contig count | 4,388 | 599 | 1,132 | 1,731 | 1,516 | 1,258 | 6,083 | 144,828 |
| Contig N50 (bp) | 133,624 | 711,969 | 41,606 | 494,779 | 448,074 | 478,824 | 78,941 | 42,786 |
| Scaffold count | 4,388 | 599 | 1,132 | 1,731 | 1,516 | 1,243 | 3,687 | 143,874 |
| Scaffold N50 (bp) | 133,624 | 711,969 | 41,606 | 494,779 | 448,074 | 484,692 | 874,516 | 54,714 |
| GC content (%) | 41.49 | 41.75 | 41.75 | 41.75 | 41.62 | 41.88 | 42.2 | 42.72 |
| TEs (bp) | 67,687,512 | 37,394,886 | 9,297,197 | 46,692,083 | 25,542,314 | 25,822,786 | 16,928,725 | 9,923,944 |
| TEs (%) | 23.47 | 22.50 | 21.51 | 22.30 | 17.41 | 17.86 | 12.36 | 7.16 |
| BUSCO (%) | | | | | | | | |
| Complete | 99.6 | 97.1 | 25.6 | 98.5 | 95.1 | 99.0 | 98.7 | 98.7 |
| Single-copy | 63.1 | 91.4 | 25.1 | 76.3 | 93.8 | 97.6 | 98.0 | 98.5 |
| Duplicated | 36.5 | 5.7 | 0.5 | 22.2 | 1.3 | 1.4 | 0.7 | 0.2 |
| Fragmented | 0.1 | 0.8 | 2.4 | 0.2 | 0.4 | 0.3 | 0.3 | 0.6 |
| Missing | 0.3 | 2.1 | 72.0 | 1.3 | 4.5 | 0.7 | 1.0 | 0.7 |

Table S2: **Statistics for A4 genome assemblies.** FALCON-Unzip_p = primary contigs; FALCON-Unzip_h = haplotigs; FALCON-Unzip_ph = primary + haplotigs.

| | Canu | Falcon-Unzip_p | Falcon-Unzip_h | Falcon-Unzip_ph | wtdbg2 | Flye | Supernova | SPAdes |
|---|---|---|---|---|---|---|---|---|
| Assembly size (bp) | 141,737,450 | 141,292,095 | 13,036,468 | 154,328,563 | 137,203,985 | 135,706,987 | 126,833,864 | 135,820,998 |
| Contig count | 181 | 107 | 289 | 396 | 431 | 234 | 3115 | 119370 |
| Contig N50 (bp) | 21,369,333 | 20,430,351 | 47,610 | 16,510,272 | 13,821,893 | 5,389,879 | 182,420 | 74,783 |
| Scaffold count | 181 | 107 | 289 | 396 | 431 | 229 | 1,818 | 118,640 |
| Scaffold N50 (bp) | 21,369,333 | 20,430,351 | 47,610 | 16,510,272 | 13,821,893 | 6,405,908 | 5,040,789 | 98,019 |
| GC content (%) | 42.07 | 42.14 | 41.89 | 42.12 | 41.83 | 42.11 | 42.27 | 42.33 |
| TEs (bp) | 21580457 | 22,804,030 | 1,803,901 | 24,607,939 | 20,832,191 | 18,259,520 | 11,339,728 | 9,393,794 |
| TEs (%) | 15.23 | 16.14 | 13.84 | 15.95 | 15.18 | 13.46 | 8.94 | 6.92 |
| BUSCO (%) | | | | | | | | |
| Complete | 99.4 | 99.2 | 8.7 | 99.4 | 94.6 | 99.5 | 99.2 | 99.2 |
| Single-copy | 98.5 | 98.7 | 8.6 | 92.5 | 94.1 | 99.1 | 98.9 | 99.1 |
| Duplicated | 0.9 | 0.5 | 0.1 | 6.9 | 0.5 | 0.4 | 0.3 | 0.1 |
| Fragmented | 0.2 | 0.2 | 0.8 | 0.2 | 0.2 | 0.2 | 0.3 | 0.3 |
| Missing | 0.4 | 0.6 | 90.5 | 0.4 | 5.2 | 0.3 | 0.5 | 0.5 |

Table S3: **Number of TEs shared between A7 and DSPR strains.** "#Overlap_A7" represents the number of TEs shared between A7 and each of the DSPR strain. Only non-*INE-1*, non-nested insertions from normal recombination regions, with flank alignment support for both sides, and for which the TAF could be calculated were included in this analysis.

| Strain | #Overlap_A7 |
|---|---|
| A1 | 8 |
| A2 | 11 |
| A3 | 7 |
| A4 | 5 |
| A5 | 15 |
| A5 | 7 |
| A7 | 658 |
| AB8 | 8 |
| B1 | 10 |
| B2 | 7 |
| B3 | 134 |
| B4 | 9 |
| B6 | 16 |

Table S4: **Feature comparison between long-read non-reference TE detection methods.** LoRTE does not predict genotypes but may flag TEs as "Possible polymorphism" if there is conflicting evidence regarding the presence/absence of a given insertion. This indicates the insertion is heterozygous or potentially a polymorphism if multiple individuals were pooled together for sequencing (Disdero and Filee, 2017).

| | TELR | LoRTE | PALMER | TLDR | xTea | rMETL | nanotei |
|---|---|---|---|---|---|---|---|
| PubMed ID | - | 28405230 | 31853540 | 33186547 | 34158502 | 30759188 | N.A. |
| Species-agnostic | Yes | Yes | Yes | Yes | No | Yes | No |
| Predicts TSD | Yes | No | Yes | Yes | Yes | No | No |
| TE sequence | Polished local assembly | Representative raw read | Representative raw read | Consensus sequence | Unpolished local assembly | Yes | No |
| Estimates TAF | Yes | No | No | No | No | No | No |
| Predicts genotype | Yes | Polymorphic TE * | No | No | No | Yes | No |
| Available in Bioconda | Yes | No | No | No | Yes | Yes | No |

Table S5: **TELR performance benchmark using genome-wide synthetic data from ISO1 and A4 genome assemblies.** Non-reference TE insertion predictions made by TELR using the A4 genome assembly as reference were evaluated against curated TE annotations in ISO1 lifted over to A4 coordinates (see Section 1.1 for details). "Zygosity" was simulated by controlling the ratio of simulated reads generated from ISO1 and A4 (see details in Section 1.1). "#True_Positives" and "#False_Positives" represent the number of predictions that match and doesn't match with curated TE annotations in ISO1 lifted over to A4 coordinates, respectively. "False Negatives" represents the number of curated TE annotations in ISO1 lifted over to A4 coordinates that were not predicted by TELR. "Precision" represents the number of true positives divided by total number of predictions made by TELR. "Recall" represents the number of true positives divided by total number of curated TE annotations in ISO1 lifted over to A4 coordinates. Only non-*INE-1*, non-nested insertions from normal recombination regions, with flank alignment support for both sides, and for which the TAF could be calculated were included in this analysis.

| Ploidy | Zygosity | Coverage | #Total_Pred | #True_Positives | #False_Positives | #False_Negatives | Precision | Recall |
|---|---|---|---|---|---|---|---|---|
| diploid | homozygous | 50 | 443 | 434 | 9 | 184 | 98.0% | 70.2% |
| diploid | homozygous | 100 | 476 | 467 | 9 | 151 | 98.1% | 75.6% |
| diploid | homozygous | 150 | 500 | 490 | 10 | 128 | 98.0% | 79.3% |
| diploid | homozygous | 200 | 489 | 480 | 9 | 138 | 98.2% | 77.7% |
| diploid | heterozygous | 50 | 363 | 358 | 5 | 260 | 98.6% | 57.9% |
| diploid | heterozygous | 100 | 461 | 450 | 11 | 168 | 97.6% | 72.8% |
| diploid | heterozygous | 150 | 478 | 469 | 9 | 149 | 98.1% | 75.9% |
| diploid | heterozygous | 200 | 472 | 462 | 10 | 156 | 97.9% | 74.8% |
| tetraploid | simplex | 50 | 116 | 116 | 0 | 502 | 100.0% | 18.8% |
| tetraploid | simplex | 100 | 366 | 361 | 5 | 257 | 98.6% | 58.4% |
| tetraploid | simplex | 150 | 432 | 425 | 7 | 193 | 98.4% | 68.8% |
| tetraploid | simplex | 200 | 452 | 446 | 6 | 172 | 98.7% | 72.2% |
| tetraploid | duplex | 50 | 357 | 352 | 5 | 266 | 98.6% | 57.0% |
| tetraploid | duplex | 100 | 445 | 437 | 8 | 181 | 98.2% | 70.7% |
| tetraploid | duplex | 150 | 480 | 472 | 8 | 146 | 98.3% | 76.4% |
| tetraploid | duplex | 200 | 481 | 472 | 9 | 146 | 98.1% | 76.4% |
| tetraploid | triplex | 50 | 431 | 423 | 8 | 195 | 98.1% | 68.4% |
| tetraploid | triplex | 100 | 486 | 477 | 9 | 141 | 98.1% | 77.2% |
| tetraploid | triplex | 150 | 472 | 462 | 10 | 156 | 97.9% | 74.8% |
| tetraploid | triplex | 200 | 474 | 465 | 9 | 153 | 98.1% | 75.2% |
| tetraploid | quadruplex | 50 | 455 | 449 | 6 | 169 | 98.7% | 72.7% |
| tetraploid | quadruplex | 100 | 475 | 465 | 10 | 153 | 97.9% | 75.2% |
| tetraploid | quadruplex | 150 | 490 | 481 | 9 | 137 | 98.2% | 77.8% |
| tetraploid | quadruplex | 200 | 480 | 469 | 11 | 149 | 97.7% | 75.9% |

Table S6: **Performance benchmark for intra-sample TE allele copy number classifier on diploid genome.** TELR predictions on synthetic data from ISO1 and A4 genome assemblies were used as input for the classifier. "Zygosity" was simulated by controlling the ratio of simulated reads generated from ISO1 and A4 (see details in Section 1.1). "Precision" represents the proportion of TE allele copy number being correctly classified. Only non-*INE-1*, non-nested insertions from normal recombination regions, with flank alignment support for both sides, and for which the TAF could be calculated were included in this analysis.

| Ploidy | Zygosity | Coverage | #Total_Pred | #TE_CN_1 | #TE_CN_2 | #Unclassified | Precision |
|---|---|---|---|---|---|---|---|
| diploid | homozygous | 50 | 443 | 0 | 443 | 0 | 100.0% |
| diploid | homozygous | 100 | 476 | 1 | 475 | 0 | 99.8% |
| diploid | homozygous | 150 | 500 | 1 | 499 | 0 | 99.8% |
| diploid | homozygous | 200 | 489 | 1 | 488 | 0 | 99.8% |
| diploid | heterozygous | 50 | 363 | 361 | 2 | 0 | 99.4% |
| diploid | heterozygous | 100 | 461 | 460 | 1 | 0 | 99.8% |
| diploid | heterozygous | 150 | 478 | 476 | 2 | 0 | 99.6% |
| diploid | heterozygous | 200 | 472 | 470 | 2 | 0 | 99.6% |

Table S7: **Performance benchmark for intra-sample TE allele copy number classifier on tetraploid genome.** TELR predictions on synthetic data from ISO1 and A4 genome assemblies were used as input for the classifier. "Zygosity" was simulated by controlling the ratio of simulated reads generated from ISO1 and A4 (see details in Section 1.1). "Precision" represents the proportion of TE allele copy number being correctly classified. Only non-*INE-1*, non-nested insertions from normal recombination regions, with flank alignment support for both sides, and for which the TAF could be calculated were included in this analysis.

| Ploidy | Zygosity | Coverage | #Total_Pred | #TE_CN_1 | #TE_CN_2 | #TE_CN_3 | #TE_CN_4 | Precision |
|---|---|---|---|---|---|---|---|---|
| tetraploid | simplex | 50 | 116 | 104 | 12 | 0 | 0 | 89.7% |
| tetraploid | simplex | 100 | 366 | 360 | 6 | 0 | 0 | 98.4% |
| tetraploid | simplex | 150 | 432 | 429 | 3 | 0 | 0 | 99.3% |
| tetraploid | simplex | 200 | 452 | 446 | 5 | 0 | 1 | 98.7% |
| tetraploid | duplex | 50 | 357 | 9 | 332 | 16 | 0 | 93.0% |
| tetraploid | duplex | 100 | 445 | 7 | 430 | 7 | 1 | 96.6% |
| tetraploid | duplex | 150 | 480 | 4 | 470 | 4 | 2 | 97.9% |
| tetraploid | duplex | 200 | 481 | 4 | 470 | 6 | 1 | 97.7% |
| tetraploid | triplex | 50 | 431 | 0 | 17 | 408 | 6 | 94.7% |
| tetraploid | triplex | 100 | 486 | 0 | 10 | 472 | 4 | 97.1% |
| tetraploid | triplex | 150 | 472 | 0 | 3 | 466 | 3 | 98.7% |
| tetraploid | triplex | 200 | 474 | 2 | 3 | 466 | 3 | 98.3% |
| tetraploid | quadruplex | 50 | 455 | 0 | 1 | 0 | 454 | 99.8% |
| tetraploid | quadruplex | 100 | 475 | 0 | 1 | 0 | 474 | 99.8% |
| tetraploid | quadruplex | 150 | 490 | 0 | 1 | 0 | 489 | 99.8% |
| tetraploid | quadruplex | 200 | 480 | 1 | 1 | 0 | 478 | 99.6% |

### 3 Supplementary Figures

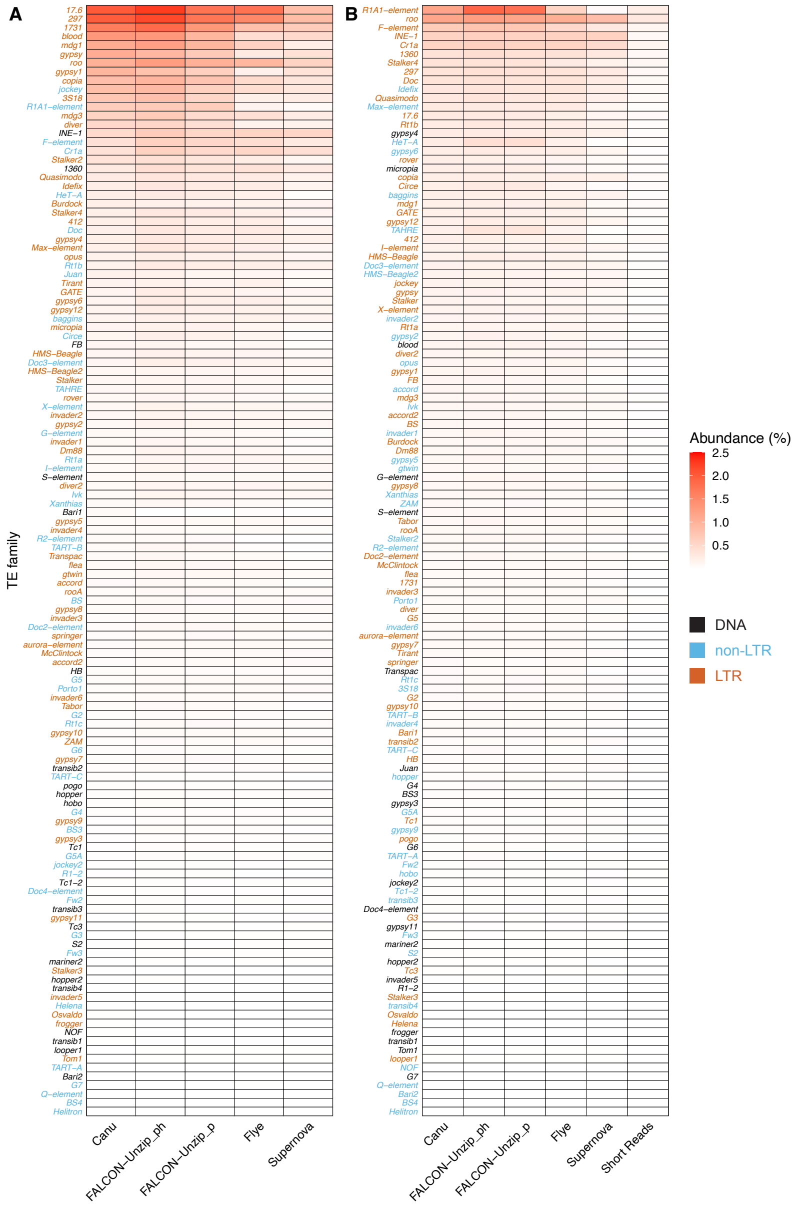


Figure S1: **TE abundance varies substantially between assembly methods.** TE abundance was estimated for different assemblies and directly from raw Illumina reads in S2R+ (A) and A4 (B) using RepeatMasker and the curated canonical *D. melanogaster* TE library. TE family names were colorized by TE type.

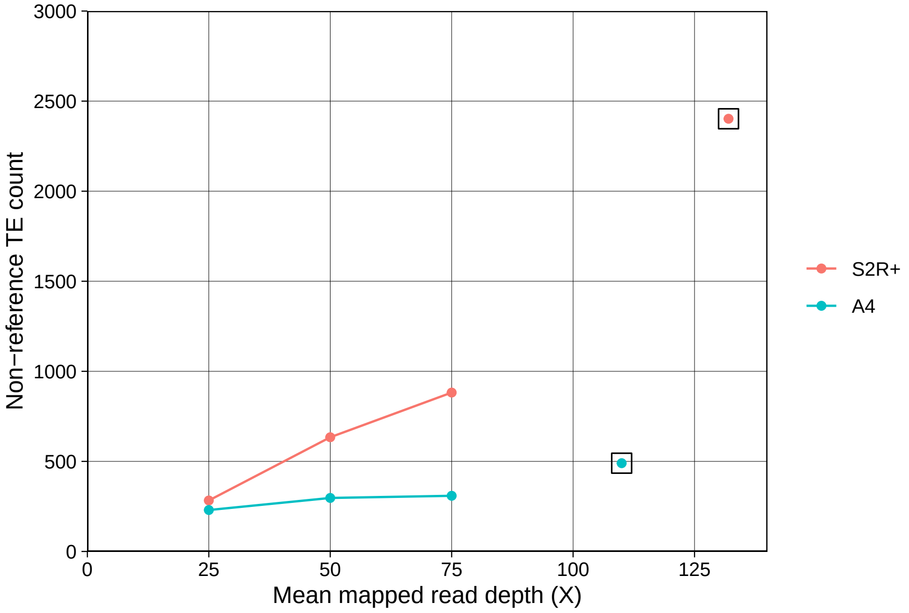


Figure S2: **Effect of mean mapped read depth in the number of non-reference TE predictions.** Comparison of the number of non-reference TEs predicted by TELR using length- and depth-normalized long-read datasets for S2R+ and A4. Reads from both datasets were split into 8kb segments and sub-sampled to achieve the desired mean mapped read depth. The boxed points indicate the number of TE predictions made by TELR when the full non-normalized data was supplied. Only non-*INE-1*, non-nested insertions from normal recombination regions, with flank alignment support for both sides, and for which the TAF could be calculated were included in this analysis.

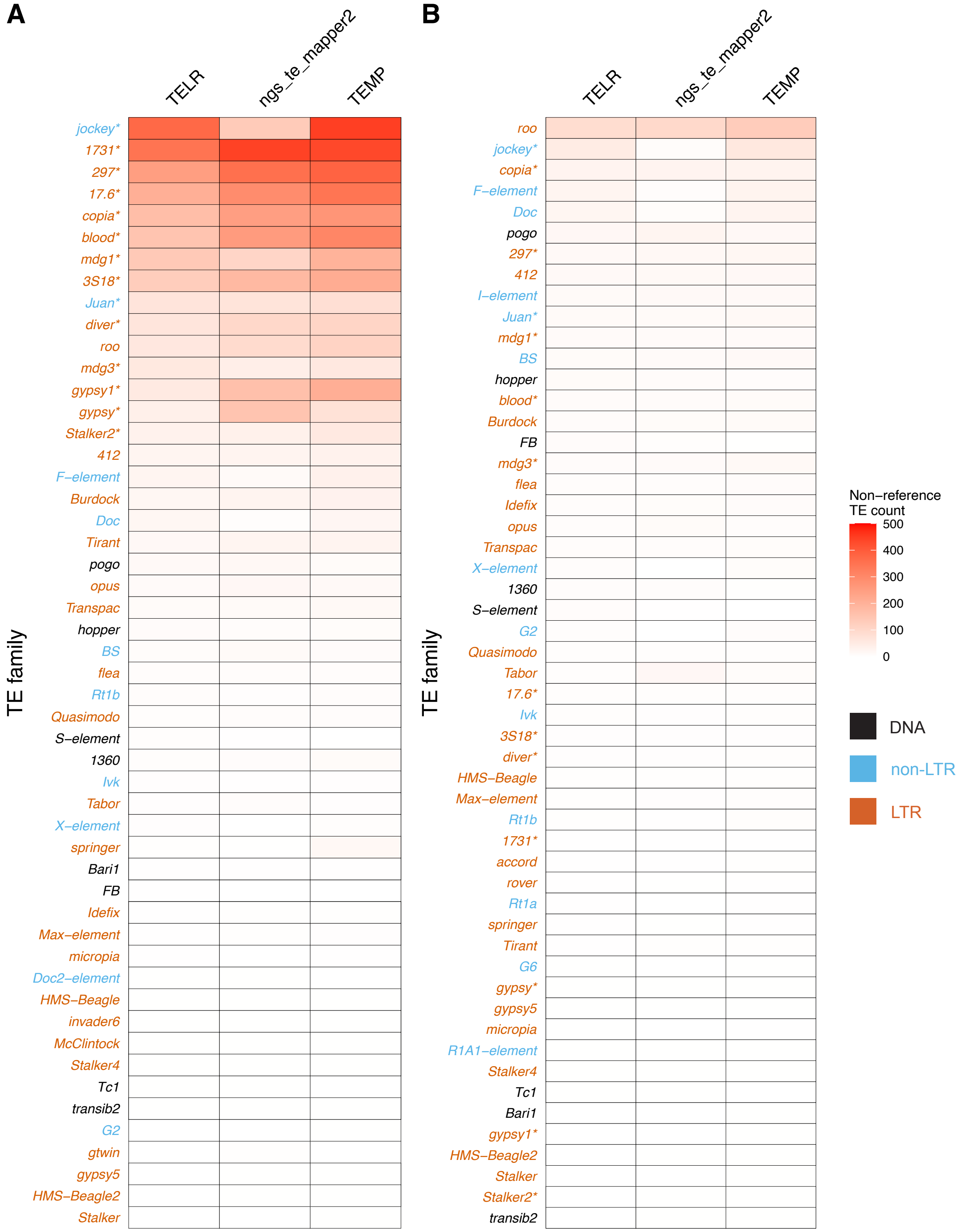


Figure S3: **Abundant TE families detected by TELR are corroborated by short-read methods.** Heatmaps represent number of non-reference TEs separated by TE families for S2R+ (A) and A4 (B). TE family names were colorized by TE type. TE families enriched in S2R+ are indicated by asterisks. Only non-*INE-1*, non-nested insertions from normal recombination regions, with flank alignment support for both sides, and for which the TAF could be calculated were included in this analysis.

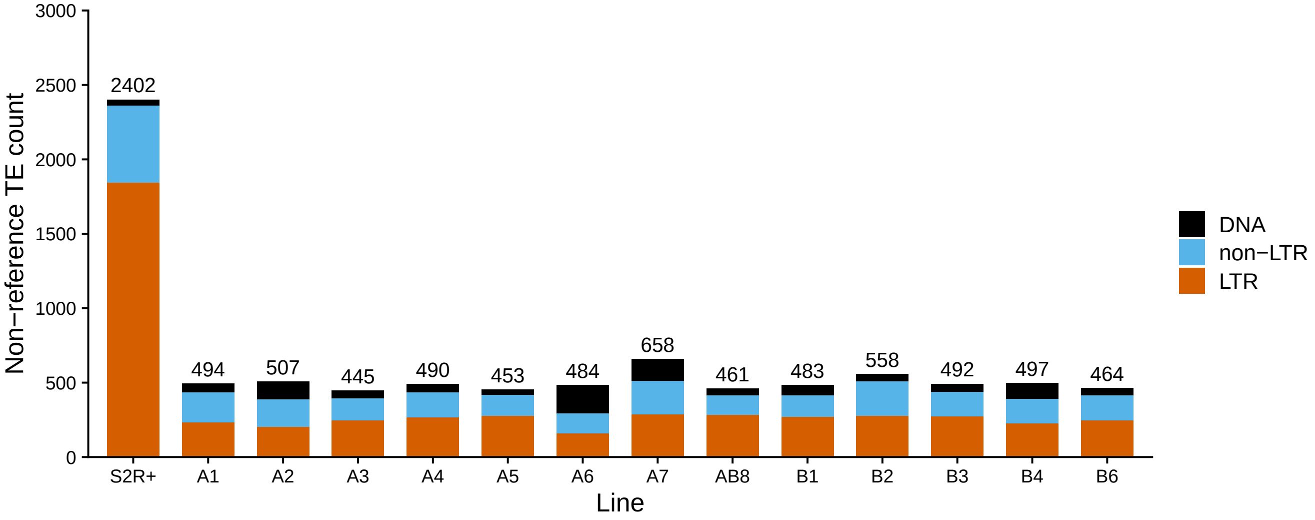


Figure S4: **Increased number of non-reference TE insertions in S2R+ compared to inbred fly stocks derived from natural populations.** Number of non-reference TE insertions predicted by TELR for S2R+ and inbred stocks from geographically diverse natural fly populations (Chakraborty *et al.*, 2019). The bars were colorized by TE type. Number on top of each bar represents total number of non-reference TE insertions for a given line. Only non-*INE-1*, non-nested insertions from normal recombination regions, with flank alignment support for both sides, and for which the TAF could be calculated were included in this analysis.

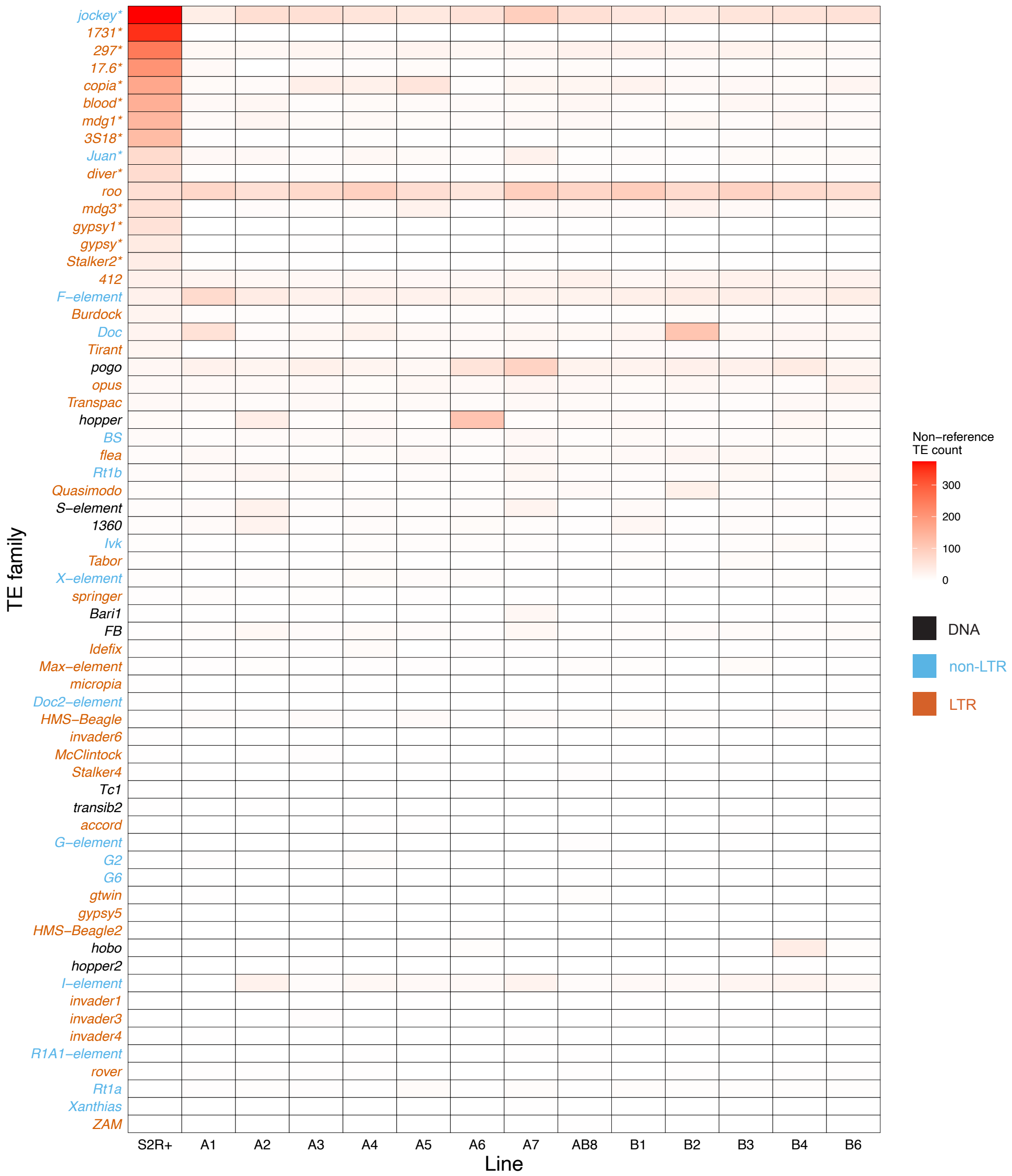


Figure S5: **Increased TE abundance and unique profile of TE activation in S2R+ compared to multiple fly strains.** Heatmap showing the copy number of non-reference insertions from multiple TE families in S2R+ and inbred fly strains from the Drosophila Synthetic Population Resource (DSPR) (King *et al.*, 2012). TE families are ordered according to their abundance in S2R+. TE family names were colorized by TE type. TE families enriched in S2R+ are indicated by asterisks. Only non-*INE-1*, non-nested insertions from normal recombination regions, with flank alignment support for both sides, and for which the TAF could be calculated were included in this analysis.

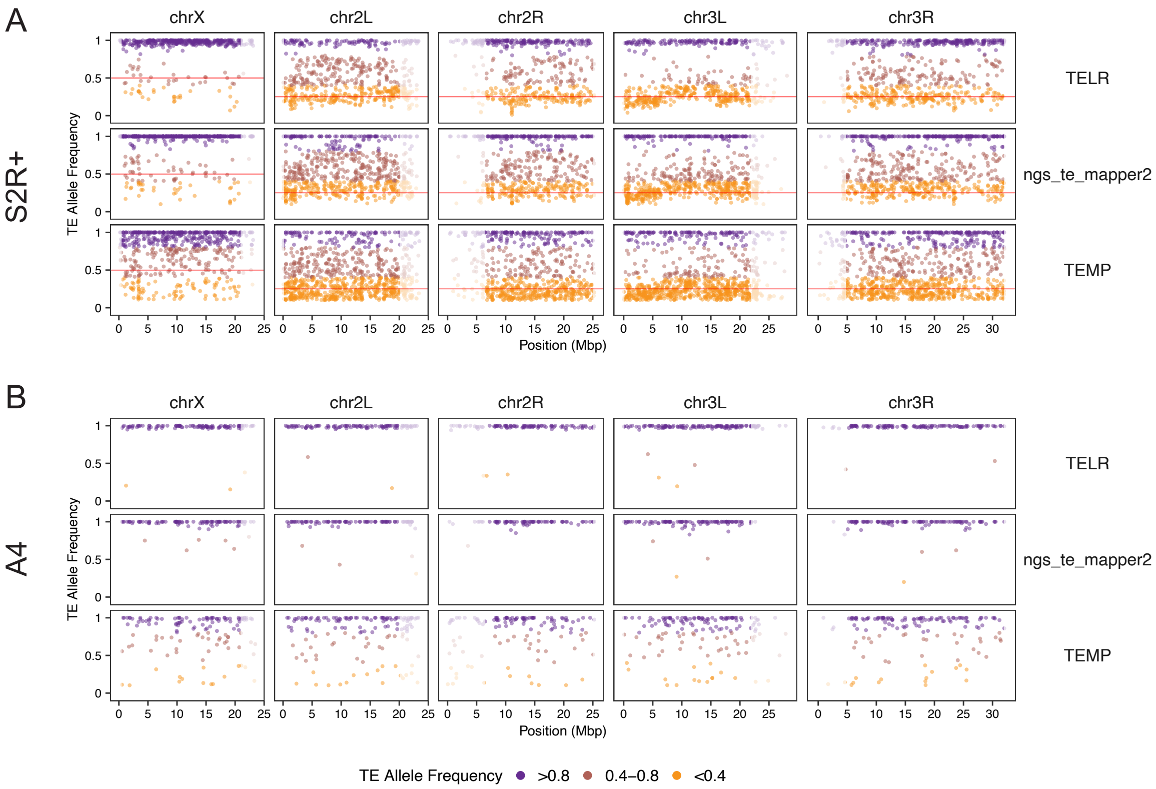


Figure S6: **Comparison of TAF genome-wide distributions between TELR and short-read methods for S2R+ and A4.** Only non-*INE-1*, non-nested insertions with flank alignment support for both sides, and for which the TAF could be calculated were included in this analysis. Low recombination regions are shaded in grey.

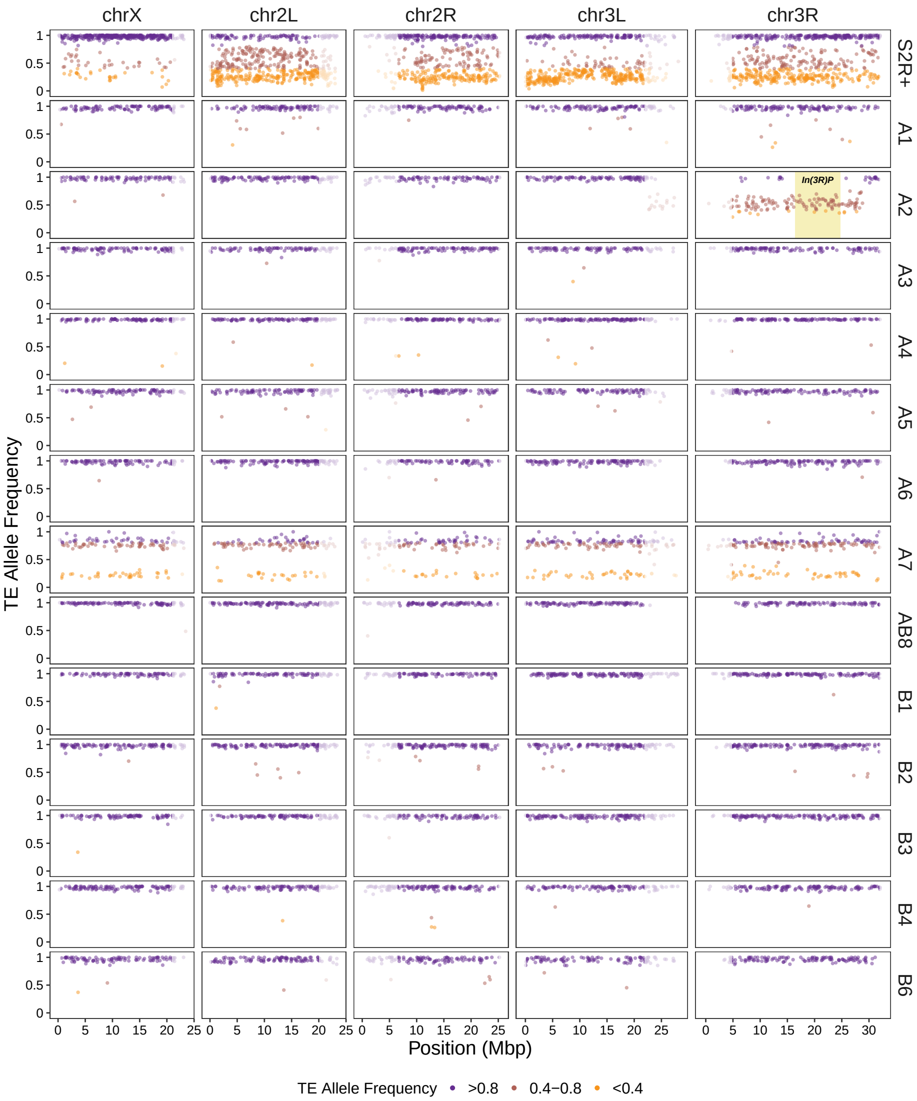


Figure S7: **TAF genome-wide distributions for S2R+ and *D. melanogaster* inbred fly strains.** Strain A2 has a known heterozygous inversion, *In(3R)P*, that prevents the complete inbreeding in chr3R (King *et al.*, 2012). Only non-*INE-1*, non-nested insertions with flank alignment support for both sides, and for which the TAF could be calculated were included in this analysis. Low recombination regions are shaded in grey.

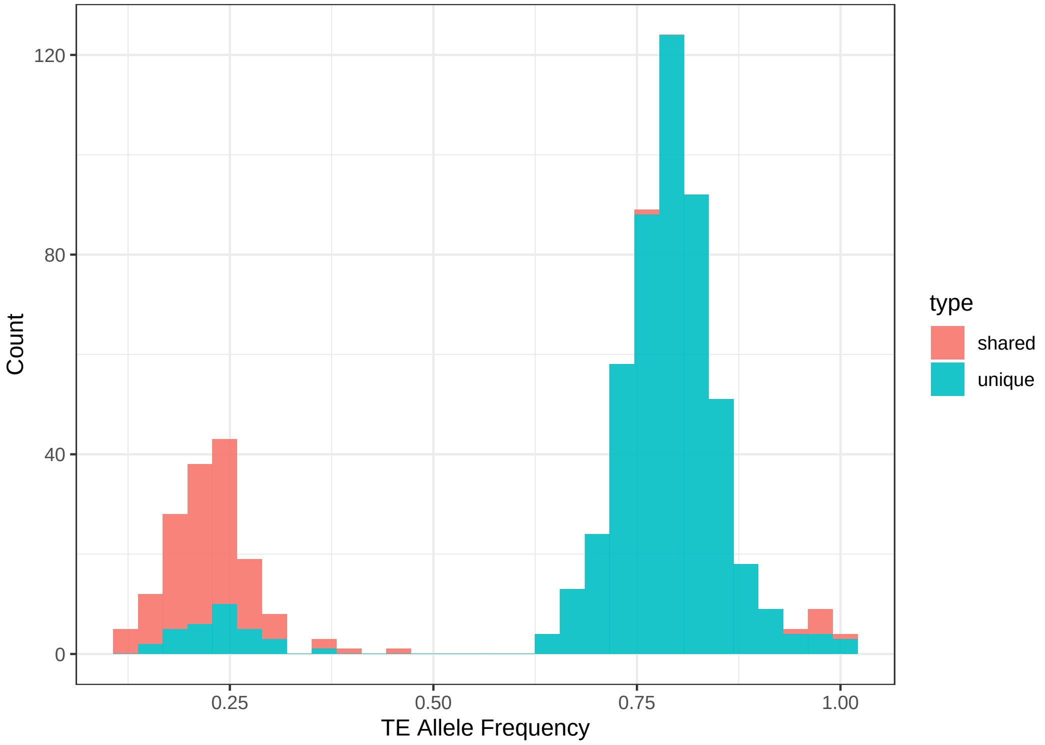


Figure S8: **TAF distribution of non-reference insertions in A7.** The histogram is colorized based on whether the TE insertion in strain A7 is shared with strain B3. Only non-*INE-1*, non-nested insertions from normal recombination regions, with flank alignment support for both sides, and for which the TAF could be calculated were included in this analysis.

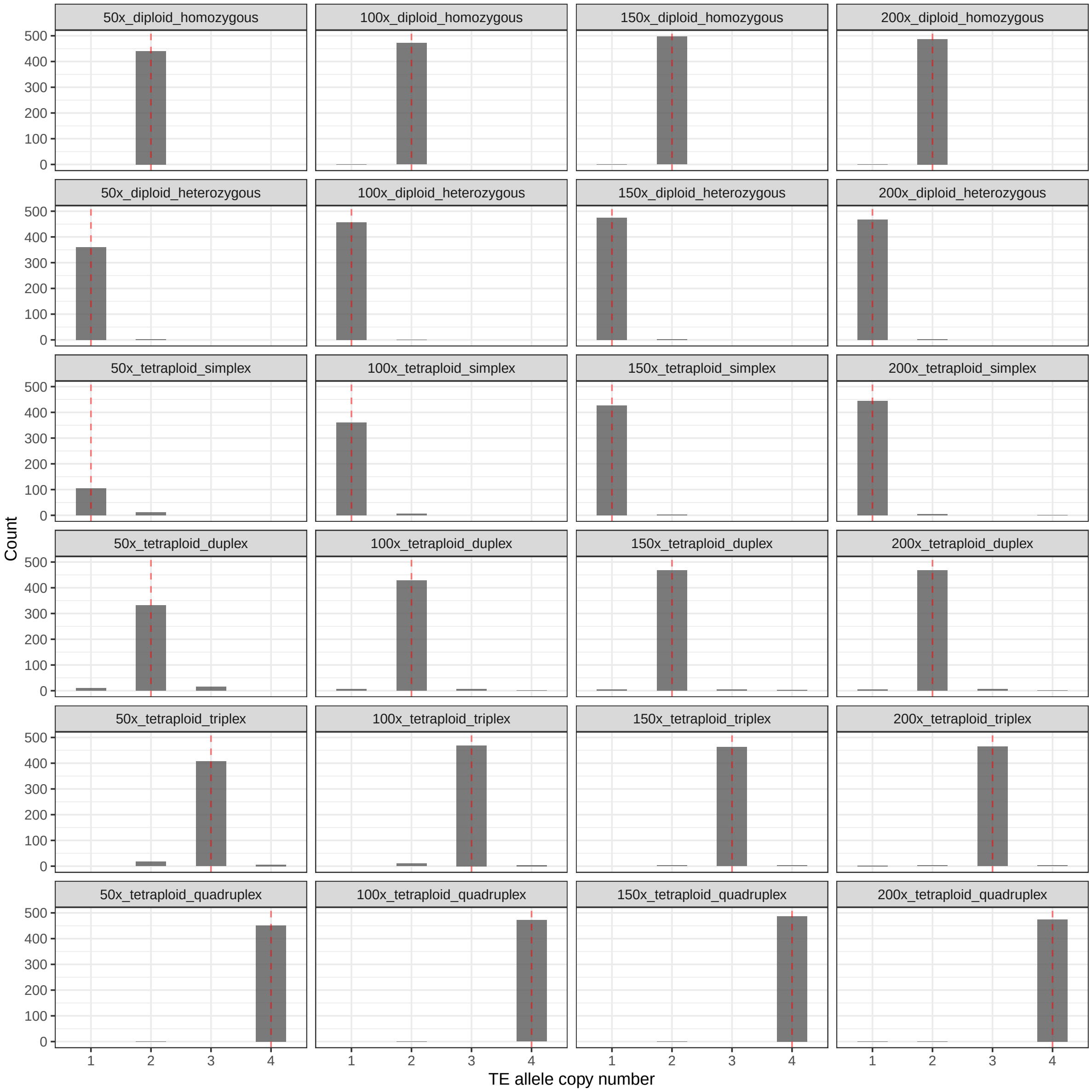


Figure S9: **TE allele copy number distribution using synthetic long read sequencing data.** Histograms of TE allele copy number on synthetic long read data under different ploidy, zygosity and coverage settings. See section 1.1 on details about how the synthetic data was generated and how the TE allele copy number was estimated. The expected TE allele copy number under a given ploidy and zygosity setting is marked in dashed red line.

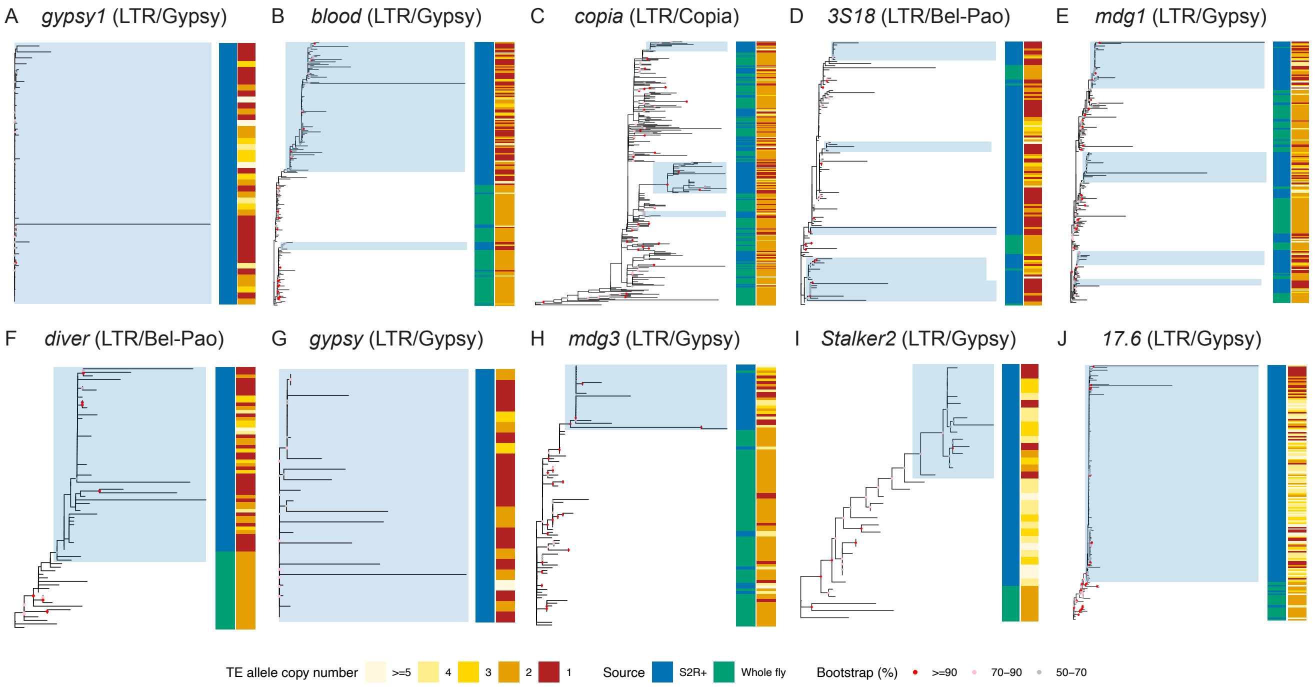


Figure S10: **Single and multiple TE source lineage activation in S2R+ cell line.** Non-reference TE insertion sequences from S2R+ and 11 inbred *Drosophila* fly strains were predicted and assembled by TELR. Full length TE sequences for each family were aligned using MAFFT (v7.487) (Katoh and Standley, 2013). The multiple sequence alignments were used as input in IQ-TREE (v2.1.4-beta) (Minh *et al.*, 2020) to build unrooted trees for *gypsy1* (A), *blood* (B), *copia* (C), *3S18* (D), *mdg1* (E), *diver* (F), *gypsy* (G), *mdg3* (H), *Stalker2* (I) and *17.6* (J) (A) elements using maximum likelihood approach. The sample source and TE copy number were annotated in the sidebars. Blue shading indicates TE expansion event in S2R+ from a single source lineage based on the following criteria: 1) All sequences should form a monophyletic clade, 2) The bootstrap support for the clade should be equal to or higher than 70%, and 3) The proportion of post-tetraploid cell-line-specific TE insertions (i.e. TE allele copy number equal to one) within the clade should be equal to or higher than 30%.

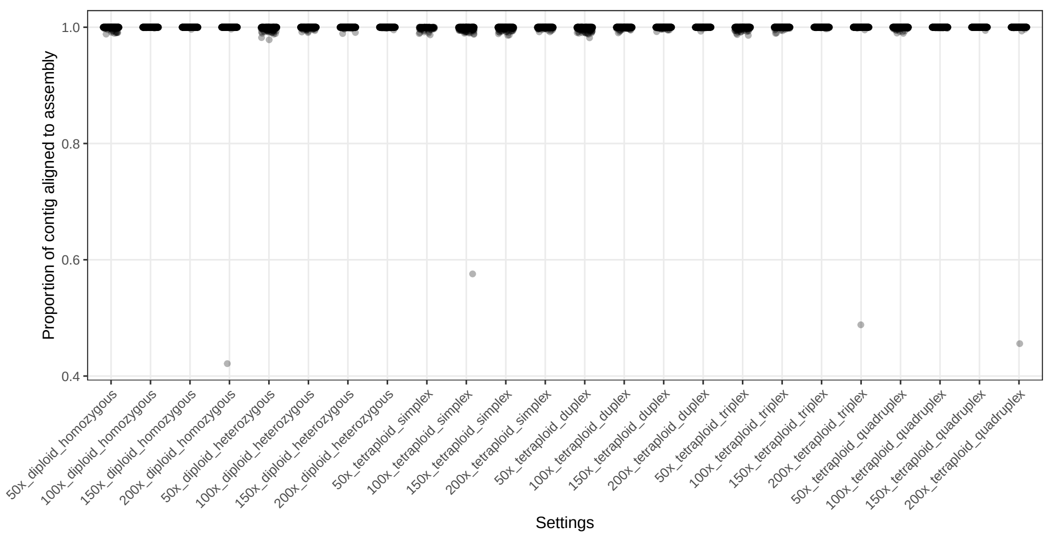


Figure S11: **Distribution on proportions of TE loci assembled by TELR aligned to the ISO1 genome assembly.** For a given TELR run using synthetic long read sequencing data generated under a specific coverage, ploidy, and zygosity setting, each TE sequence plus 500bp flanking sequences on 5' and 3' side of the TE locus in the local contig assembly made by TELR (referred to as "TELR TE locus") was aligned to the ISO1 genome assembly. "Proportion of contig aligned to assembly" on the y-axis represents the proportion of each TELR TE locus that can be aligned to ISO1. See details in Section 1.3.

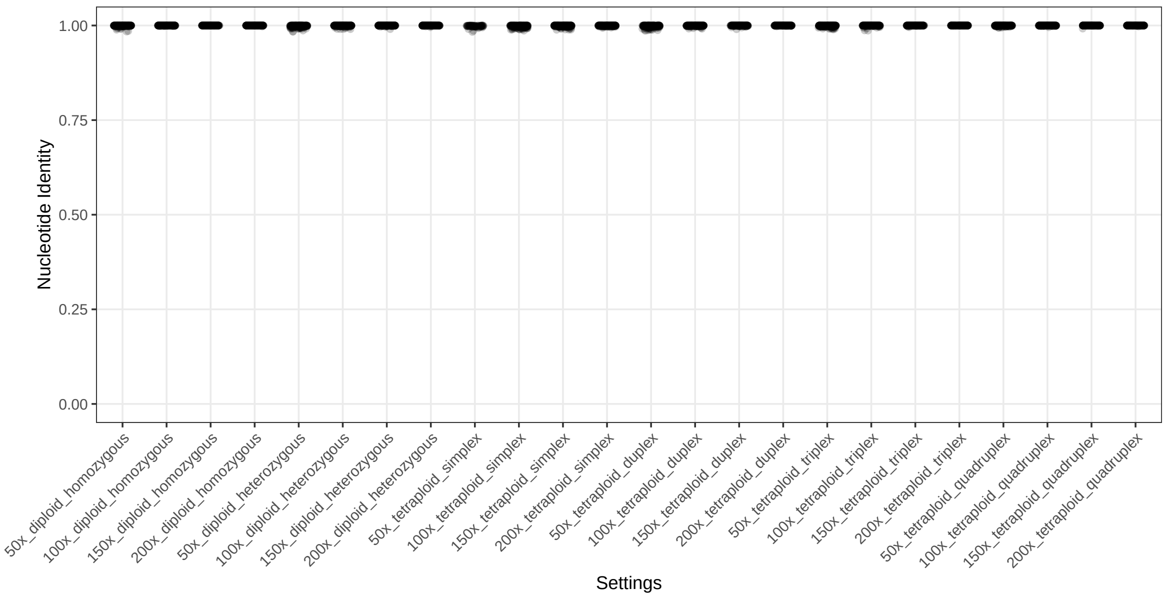


Figure S12: **Distribution of nucleotide identity between TELR TE sequences and corresponding TE sequences in ISO1 using synthetic long read sequencing data.** For a given TELR run using synthetic long read sequencing data generated under a specific coverage, ploidy, and zygosity setting, each predicted TE sequence plus 500bp flanking sequences on 5' and 3' side of the TE locus in the local contig assembly (referred to as "TELR TE locus") was aligned to the ISO1 genome assembly. For each TE insertion predicted by TELR that can be matched with a corresponding curated TE annotation in ISO1 (FlyBase release 6.38), we aligned TELR TE sequence with TE sequences in ISO1 based on curated TE annotation using minimap2 (Li, 2018) and calculated nucleotide identity. See details in Section 1.3.

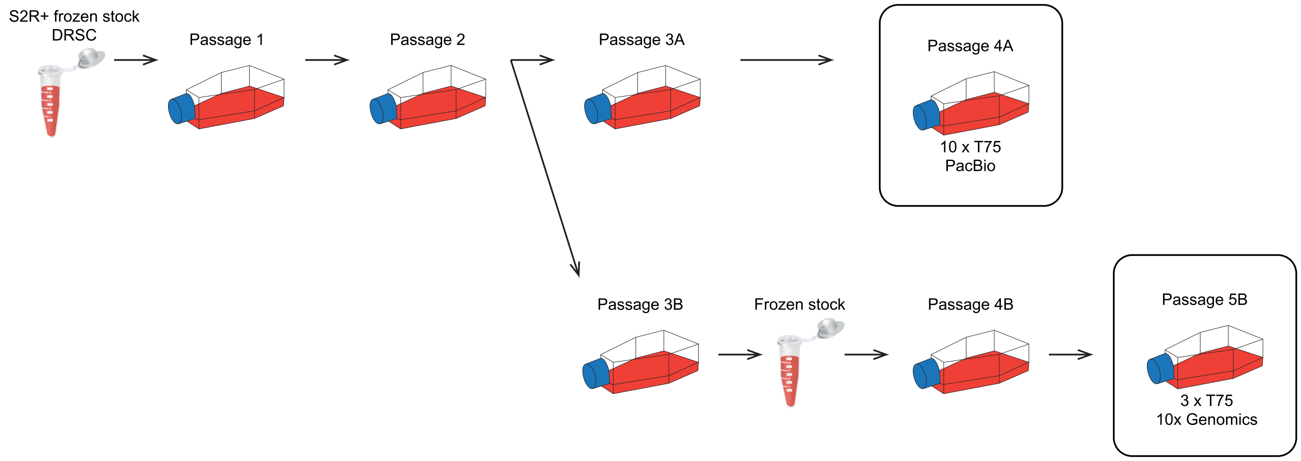


Figure S13: **Provenance of the S2R+ cells used in this study.** Cells were harvested from passages 4A and 5B for PacBio and 10x Genomics linked-read DNA sequencing, respectively.
